# Neuromodulation Exerts Feedback and Feedforward Control of Action Selection

**DOI:** 10.1101/2020.06.08.140210

**Authors:** Fengqiu Diao, Nathan Peabody, Benjamin H. White

## Abstract

To be effective, behavioral choices must result in actions that are appropriate to an animal’s needs and environmental circumstances. In addition, the actions must be ones the animal can sustain until its needs are met. This aligning of goals, action, and motivation requires the coordinated activity of multiple brain circuits, but how such coordination is achieved is poorly understood. Here, we show how the insect hormone Bursicon coordinates the selection and sustains execution of a behavior in newly emerged adult *Drosophila.* Such flies must expand and harden their wings after metamorphosis, but they choose to delay expansion in confined conditions. We show that the decision to expand is mediated by an environmentally-sensitive, positive feedback loop in which Bursicon promotes its own sustained release. Released Bursicon then modulates motor neurons to promote wing expansion behavior. Bursicon thus exerts feedforward and feedback control to coordinately select and motivate a goal-directed action.

## Introduction

Physiological needs for such things as food or water are often communicated to the brain by hormones, which act on interoceptive neurons to induce persistent need states, such as hunger or thirst. Such states organize the behavioral activities of the animal to meet the required need. A common organization involves behaviors for searching the environment to identify the needed resource (e.g. food, water) followed by behaviors for exploiting the resource (e.g. eating or drinking). This division of behavior into flexible, “appetitive” search strategies followed by more stereotyped, genetically programmed, “consummatory” motor programs was first noted by the early ethologist Wallace Craig and later elaborated by Konrad Lorenz (Craig, 1996; Lorenz, 1981). Although much has been subsequently learned about the neurons and circuits that generate appetitive and consummatory motor patterns, the neural mechanisms that temporally organize these activities and adjust them to changing environmental conditions remain largely unknown.

Hormones were among the first factors proposed to help establish excitatory states in the nervous system to organize an animal’s activities towards specific goals (Beach, 1942; Tinbergen, 1951). Together with neuromodulators, which are often prominent signaling components of interoceptive neurons, hormones can act broadly within the nervous system and thus influence the activity of circuits governing distinct processes. Oxytocin, for example, both modulates the sensory salience of a pup’s cries to its mother (Marlin et al., 2015) and facilitates maternal aggression towards intruders (Bosch, 2013), and the Neuropeptide Y (NPY) homologue in insects, NPF, both promotes feeding (Chung et al., 2017) and gates the expression of olfactory memories associated with food in response to hunger (Krashes et al., 2009). Such examples, and the evolutionarily conserved association NPY with feeding (Fadda et al., 2019; Soengas et al., 2018; Tachibana and Tsutsui, 2016; Taghert and Nitabach, 2012) and oxytocin with affiliative behaviors (Grinevich and Stoop, 2018; Lefevre and Sirigu, 2017; Mitre et al., 2016), are consistent with the idea that neuromodulators can broadly organize an animal’s behavior to meet specific needs.

The mechanisms underlying such organization, however, have been difficult to elucidate because of the often subtle effects of neuromodulation and the wide scope of neuromodulator action, especially in vertebrate nervous systems (Kelly and Goodson, 2014). Nervous systems of more tractable size, such as that of the fruit fly, *Drosophila melanogaster*, where the sources and sites of action of particular neuromodulators can be determined at cellular resolution, offer advantages in this regard. A particularly useful model for understanding how hormones and neuromodulators organize behavior is the circuit that mediates ecdysis, the process by which insects shed their exoskeletons and expand a new one (Truman, 2005; Zitnan and Adams, 2012).

In the fly, the peripherally-released Ecdysis Triggering Hormone (ETH) initiates a motor program that drives emergence of the adult followed by appetitive environmental exploration in search of a suitable perch (Diao et al., 2015; White and Ewer, 2014). After perching, interoceptive neurons responsive to ETH release the neuromodulatory factor Bursicon to regulate a consummatory motor program that expands—i.e. opens and flattens—the pleated wings so that they can be hardened for flight (Dewey et al., 2004; Luan et al., 2006a). Release of Bursicon is sensitive to environmental conditions, which necessarily interposes a decision into the wing expansion process: Flies must decide either to persist in environmental search, or to perch and expand (Peabody et al., 2009). There is urgency in this decision as delaying expansion for too long results in desiccation and death. In confined—and thus adverse—environments, flies will substantially prolong environmental search to escape confinement before finally expanding their wings.

If environmental search fails to result in escape from confinement, flies will ultimately perch and expand. They will not do so, however, if a pair of Bursicon-secreting neurons in the subesophageal zone called the B_SEG_ is electrically suppressed (Luan et al., 2012). B_SEG_ activation is sufficient to terminate search and force wing expansion in confined flies, indicating that these neurons are critical for promoting the decision to expand. Interestingly, if the decision is simplified by placing flies in a spacious container and leaving them completely unperturbed, most flies with suppressed B_SEG_ neurons will nevertheless expand, although they do so using alternative motor patterns to those normally used. This suggests that the B_SEG_ likely promote expansion by affecting both motor program execution and composition. They are thus particularly attractive candidates for unraveling how a neuromodulator might coordinately regulate motor, motivational, and decision-making circuit nodes to bias behavior towards a particular goal.

Using the Trojan exon method (Diao et al., 2015) to target and manipulate the function of neurons that express the Bursicon receptor, we here identify the downstream targets of the B_SEG_. We find that these targets divide into functionally distinct populations, which together facilitate the wing expansion decision and direct the assembly of the behaviors required to execute it. They do so by participating in either feedforward motor circuits for abdominal contraction and air swallowing—motor patterns required for wing expansion—or a feedback modulatory circuit that promotes Bursicon release from the B_SEG_ to initiate and maintain the execution of those motor patterns. By coordinately facilitating the wing expansion decision and the actions required to complete it, Bursicon neuromodulation thus directly couples the assembly, selection, and sustained execution of an ethologically important action.

## Results

### Positive feedback controls Bursicon release to motivate wing expansion

To identify and characterize neurons targeted by Bursicon released from the B_SEG_, we used the rk^pan^-Gal4 driver line (Diao and White, 2012), which expresses the transcription factor Gal4 selectively in cells that express the Bursicon receptor, Rickets (RK, Fig. 1A-A”). The group of RK-expressing neurons (hereafter referred to as RK neurons) includes neither the B_SEG_, which release Bursicon into the central nervous system (CNS) to facilitate wing expansion behavior, nor the B_AG_, which release Bursicon into the hemolymph (Fig. 1A, B). We first asked whether activation of RK neurons is able to rapidly induce wing expansion in newly emerged flies, as has been previously demonstrated for the B_SEG_. To do so, we used the heat-dependent effector, UAS-dTrpA1 and confined flies in “minichambers,” which are approximately three fly-lengths in size and have previously been shown to significantly promote appetitive search behavior and delay the initiation of wing expansion (Peabody et al., 2009). We found that activating RK neurons in confined flies shortly after eclosion immediately terminated search and rapidly induced the consummatory act of wing expansion, which includes motor patterns for abdominal contraction, proboscis extension, and cibarial pumping (Fig. 1C-E; Video 1). These motor patterns promote swallowing to fill the gut with a bubble of air, which when compressed by tonic abdominal contraction drives hemolymph into the thorax and wings to expand them. In contrast to experimental animals, control flies lacking either the UAS-dTrpA1 transgene or the rk^pan^-Gal4 driver expanded with characteristic confinement-associated delays and actively sought to escape the minichamber by walking and probing its corners (i.e. “digging”), as has been previously described (Fig. 1C, Peabody et al., 2009). Bursicon binding to RK is known to stimulate cAMP production (Luo et al., 2005; Mendive et al., 2005), and acutely increasing cAMP levels in RK neurons using the photoactivatable adenylyl cyclase, UAS-euPACα (Schroder-Lang et al., 2007), likewise abbreviated search behavior and accelerated wing expansion in confined flies (Fig. 1— figure supplement 1). These results demonstrate that activation of neurons downstream of Bursicon is sufficient to induce wing expansion, similar to the activation of the B_SEG_.

**Figure 1.**
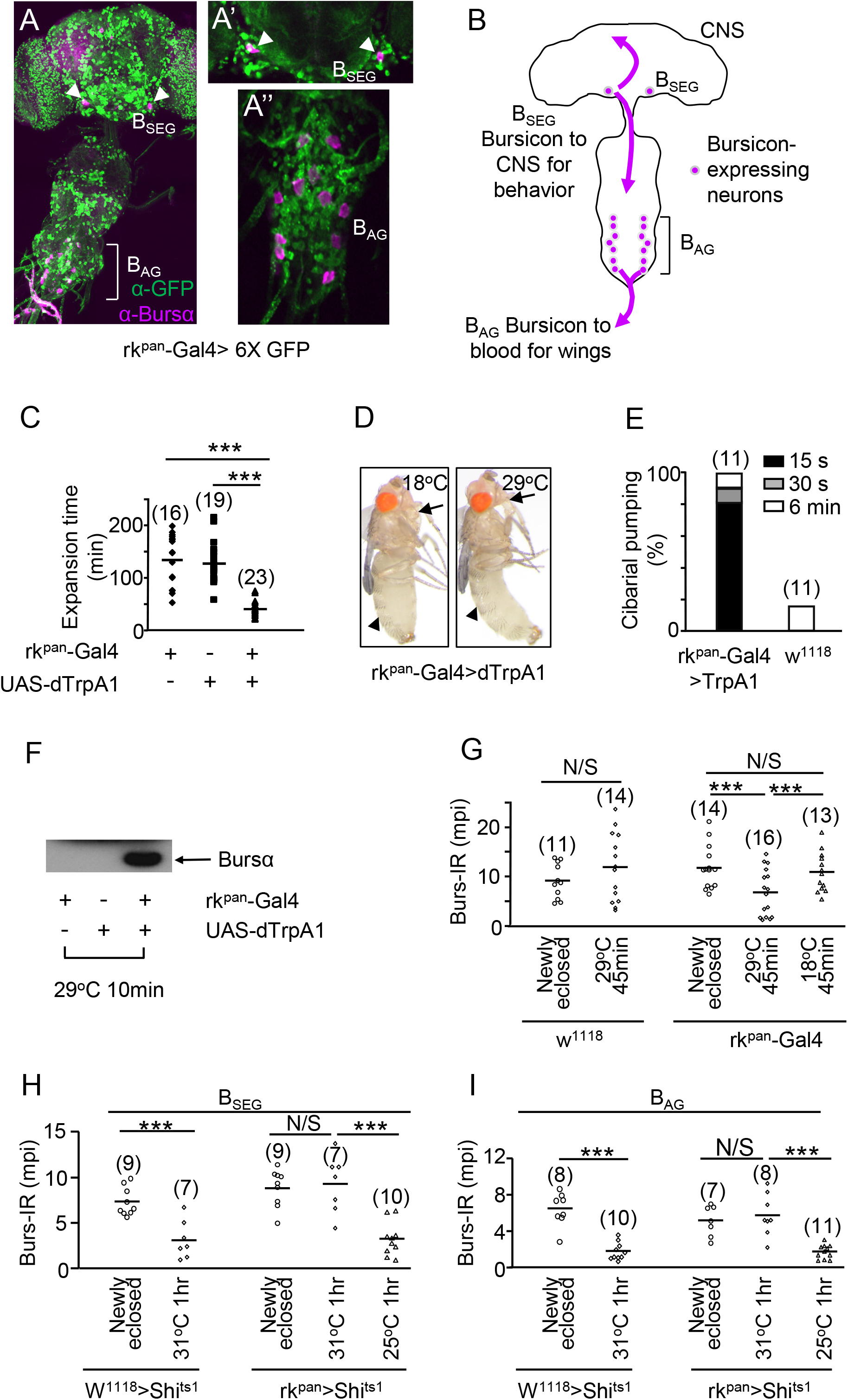
RK neurons regulate Bursicon release. **(A-A”)** Confocal image from a CNS wholemount prepared from a newly-eclosed animal expressing UAS-6XGFP under the control of the rk^pan^-Gal4 driver. To maximize detection of GFP expression, the preparation was labeled with an anti-GFP antibody (green). The preparation was also labeled with an anti-Bursα antibody to visualize the two populations of Bursicon-expressing neurons (magenta), the B_SEG_ (arrowheads) and B_AG_ (bracket). No co-labeling of the Bursicon-expressing neurons was observed in individual z-planes that included the B_SEG_ (**A’**, magenta, arrowheads) or B_AG_ (**A”**, magenta) neurons. **(B)** Schematic illustrates the distinct functional roles of Bursicon derived from the B_SEG_ and the B_AG_. B_SEG_-derived Bursicon acts within the CNS to modulate behavior; B_AG_-derived Bursicon acts hormonally on the wing epidermis. **(C)** Stimulation of RK neurons using the heat-sensitive ion channel UAS-dTrpA1 induces rapid wing expansion in confined flies. Experimental flies (rk^pan^-Gal4>dTrpA1) in minichambers expand significantly more quickly at the activating temperature (29°C), than control flies. Points represent expansion times of all individual flies of each genotype; n indicated in parentheses; bars, mean expansion times; ***, p<0.005 by unpaired, two-tailed t-test. **(D)** Motor patterns immediately induced by RK neuron activation include both abdominal contraction (arrowheads) and proboscis extension (arrows), as illustrated by comparing flies before (18°C) and during (29°C) dTrpA1 activation. **(E)** Cibarial pumping is induced by RK neuron stimulation in 82% of flies within 15 s of dTrpA1 activation (black), in 91% of flies within 30 s (light gray), and in all flies within 6 min (white). Only 18% of control flies (UAS-dTrpA1 only) display any cibarial pumping in this time. **(F)** RK neuron stimulation (10 min) induces release of Bursicon into the hemolymph from B_AG_ neurons, as determined by Western blot using an anti-Bursα antibody. 1 μL hemolymph loaded. **(G)** RK neuron stimulation (45 min) causes Bursicon secretion from the B_SEG_. Scatter plots summarizing mean pixel intensities (mpi) of B_SEG_ fiber Burs-IR from multiple animals (n in parentheses; bars indicate mean; **, p<0.01 and N/S, no significant difference by unpaired, two-tailed t-test). Levels of Burs-IR observed in newly eclosed animals of both genotypes are not significantly changed by 45 min of heat (29°C) in control animals (w^1118^) or by 45 min without heat (18°C) in experimental (rk^pan^-Gal4) animals. They are, however, significantly reduced in experimental animals in which RK neurons have been stimulated (29°C) for 45 min. **(H-I)** Suppression of synaptic transmission in RK neurons of unconfined flies using UAS-Shi^ts1^ blocks wing expansion and Bursicon release from both the B_SEG_ and B_AG_. Burs-IR in fibers of the B_SEG_ (**H**) or B_AG_ (**I**) is not reduced in animals subjected to 1 hr RK neuron suppression (31°C), but is reduced in animals spared from suppression (25°C), demonstrating that synaptic block of RK neurons inhibits Bursicon secretion. Scatter plots summarize mean pixel intensities (mpi) of Burs-IR from multiple preparations (n in parentheses; bars indicate mean; ***, p<0.005 and N/S, no significant difference by unpaired, two tailed t-test).

Activation of the B_SEG_, in addition to inducing the behaviors normally required to expand the wings, also promotes release of Bursicon into the hemolymph from the neuroendocrine B_AG_ neurons (Luan et al., 2012). The blood-borne hormone acts to plasticize the cuticle of the wings and the thorax so that they can be expanded in response to the behaviorally evoked increase in hemolymph pressure also induced by the B_SEG_. After expansion, blood-borne Bursicon secondarily induces sclerotization and pigmentation (i.e. “tanning”) of the wing and exoskeleton. We find that activation of RK neurons using UAS-dTrpA1, like activation of the B_SEG,_ induces Bursicon release into the hemolymph (Fig. 1F).

Surprisingly, we also observe Bursicon release from the B_SEG_ in response to activation of the RK neurons, as determined by the depletion of Bursicon immunoreactivity (Burs-IR) from central B_SEG_ fibers (Fig. 1G; Fig. 1—figure supplement 2A, B). This result indicates that the B_SEG_ lie downstream of neurons that they target. To rule out the possibility that low-level expression of rk^pan^-Gal4—and therefore dTrpA1— in the B_SEG_ was causing the observed loss of Bursicon immunoreactivity, we examined the ability of RK neuron activation to induce wing expansion when Gal4 activity was blocked in Bursicon-expressing neurons by the Gal4 inhibitor, Gal80. We found that wing expansion was not substantially altered by this manipulation, confirming that rk^pan^-Gal4>UAS-dTrpA1 induces expansion by activating downstream targets of the B_SEG_ (Fig. 1—figure supplement 3).

To determine whether Bursicon release from the B_SEG_ and B_AG_ is contingent on upstream activity in RK neurons, we acutely suppressed neurotransmission in these neurons in newly eclosed flies using the temperature-sensitive synaptic blocker, UAS-Shi^ts1^. This manipulation, which was performed without environmental confinement so that wing expansion was not otherwise suppressed, showed significant attenuation of Bursicon release from both the central axons of the B_SEG_ and peripheral arbors of the B_AG_ (Fig. 1H, I; Fig. 1—figure supplement 2A, C). All experimental flies at 31°C (n=87) also failed to expand their wings and did not execute wing expansion behaviors during the three hours of observation, whereas all control flies (n=174) expanded their wings normally. Taken together, these results demonstrate that RK neurons are essential components of the wing expansion circuitry and are necessary for release of the RK ligand, Bursicon, from the B_SEG_ and B_AG_. This suggests that Bursicon release is maintained by positive feedback from a circuit that includes their immediately downstream targets, the RK neurons, providing a neuronal substrate for the sustained behavioral state required for completion of the wing expansion program.

### RK neuron stimulation independently regulates behavior and Bursicon release

While it is clear that RK neurons facilitate Bursicon release, the time-locked induction of expansional behaviors that we observe upon activating these neurons is likely to be independent of Bursicon, given that Bursicon’s effect on wing expansion behavior normally proceeds over a timescale of approximately 10 min (Peabody et al., 2009). To directly examine the timecourse of Bursicon’s effects, we stimulated the B_SEG_ with UAS-dTrpA1 and compared the effects with those of stimulating the RK neurons. We found that stimuli of longer duration (>5 min) were effective in inducing wing expansion in the minichamber in both cases (Fig. 2A, B), but that the behavioral onset of this process differed considerably. For example, stimulation of the B_SEG_ for 10 min induced wing expansion behaviors only after an average delay of 7.8 min (±2.2 min, stdv), whereas stimulation of RK neurons for the same duration caused all flies to immediately perch and initiate abdominal contraction.

**Figure 2.**
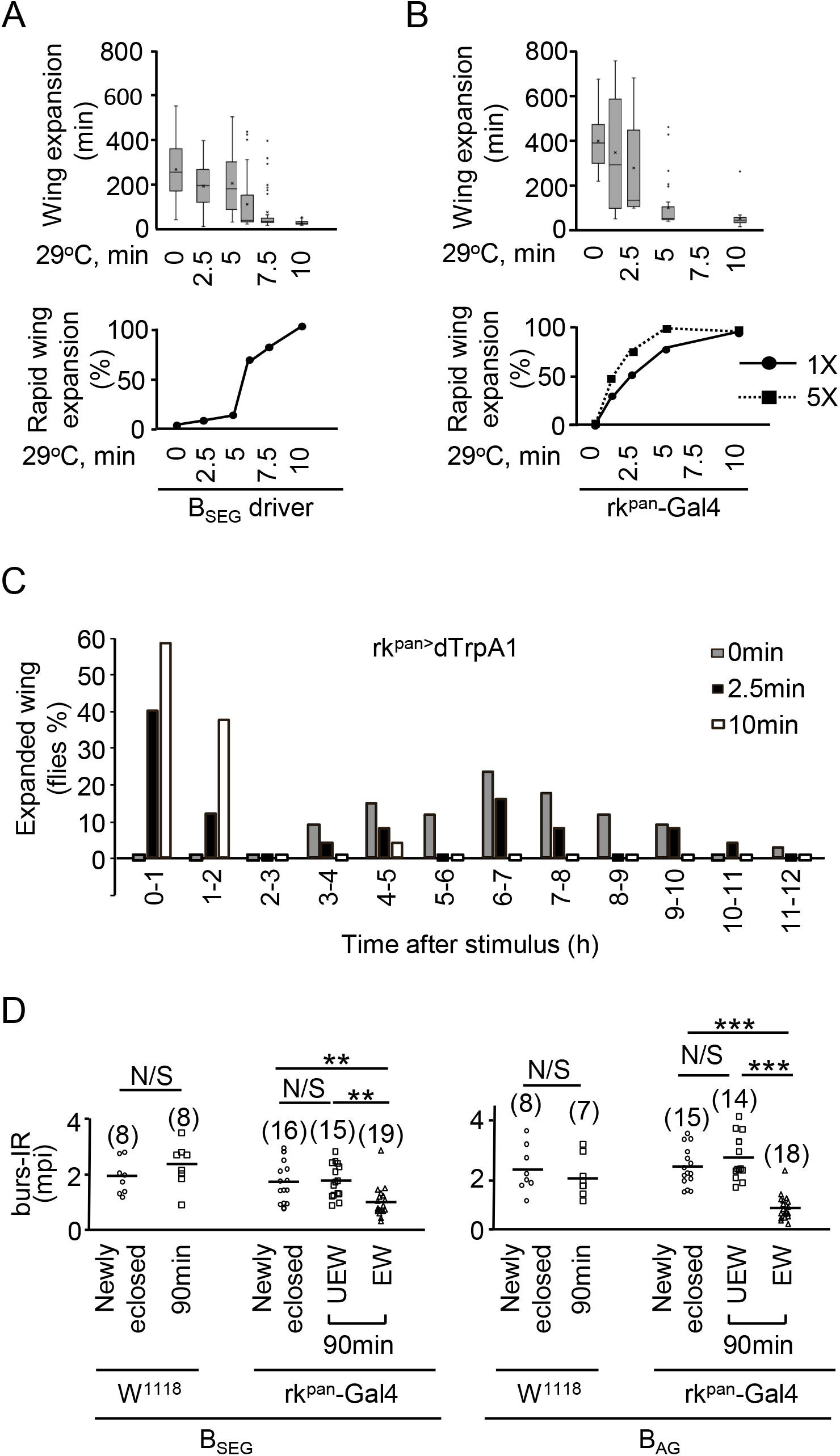
Stimulation of RK, but not B_SEG_, neurons evokes two, temporally distinct behavioral responses. (**A-B**) Effects on wing expansion of stimulating the B_SEG_ (**A**) or RK neurons (**B**) for times of 0-10 min. Box plots (upper panels) show the distribution of wing expansion times as a function of stimulus duration for stimuli of 0, 1.25, 2.5, 5, 7.5, and 10 min (and 6 min, but not 1.25 min for the B_SEG_). Newly eclosed flies were confined in the minichamber (All outliers are shown separately as dots; X indicates the mean. Graphs (lower panels) show the percentage of flies expanding rapidly in response to each stimulus. The “5X” graph shows the results of RK neuron stimulation carried out on flies in an enlarged minichamber, five times the normal length. (C) Histogram showing the percentage of flies that expand their wings during the indicated interval after stimulation of the RK neurons. A 10 min stimulus (white) induces expansion in almost all flies within 2 hr (n=24) compared with unstimulated flies (0 min, gray; n=34), which typically expand only after 4 hr. Flies receiving 2.5 min stimulation (black; n=25) divide into disjoint groups, with half showing induced expansion, and half expanding with a distribution similar to unstimulated flies. (D) Rapid wing expansion induced by 2.5 min of RK neuron stimulation is accompanied by Bursicon release from both the B_SEG_ (left panels) and B_AG_ (right panels). Scatterplots show the mean pixel intensities of Burs-IR in control (w^1118^>UAS-dTrpA1) and experimental (rk^pan^-Gal4>UAS-dTrpA1) flies directly after eclosion (Newly eclosed) or 90 min after delivery of the 2.5 min heat stimulus (29°C). Experimental flies were divided according to whether their wings expanded (EW) or remained unexpanded (UEW) after the stimulus. None of the control flies had expanded their wings within 90 min. Bursicon levels were significantly reduced in the B_SEG_ and B_AG_ projections only of experimental flies that had expanded their wings. **, p < 0.01 by unpaired, two tailed t-test; otherwise symbols as defined in Fig. 1H, I.

These results suggest that in addition to inducing Bursicon release from the B_SEG_, RK neuron stimulation independently activates downstream motor effectors of Bursicon. This was obvious at shorter stimuli (<5 min), where the immediate effects on behavior became dissociated from the induction of expansion. For example, all flies subjected to RK neuron stimulation for 2.5 min immediately initiated wing expansion behavior during the delivery of the stimulus and returned to searching behavior after stimulus termination (Video 2), but half of them re-initiated a second, successful round of expansion behavior after a brief delay of 13.0 min (± 6.7 min, stdv; n=13; Fig. 2B). Flies that did not exhibit accelerated wing expansion reverted to the search motor program on average for 363.6 min (±127.0 min, stdv; n=12), a delay similar to that seen in control animals. Flies subjected to 2.5 min stimulation thus distributed into two groups (Fig. 2C), one with long expansion times characteristic of unstimulated flies (0 min) and another with short expansion times flies characteristic of fully stimulated flies (10 min). Examination of Burs-IR in the B_SEG_ and B_AG_ fibers of these two groups 90 min after stimulus delivery revealed a strong correlation between the induction of wing expansion and the release of the hormone (Fig. 2D). Animals that expanded showed significant depletion of Burs-IR from sets of fibers, consistent with their release of Bursicon, whereas animals that remained unexpanded—and in which wing expansion had therefore not been induced—had Burs-IR levels indistinguishable from those measured in control animals.

These results show that stimulation of the RK neurons elicits two behavioral responses with distinct timecourses, a time-locked induction of wing expansion behavior that does not correlate with Bursicon release, and a somewhat delayed, but sustained induction of the full wing expansion program that does. Only delayed, sustained induction of expansion is seen in response to stimulation of the B_SEG_. We conclude that RK neurons govern two distinct types of processes: one motor, to produce the behaviors required for wing expansion, and one motivational, to promote wing expansion by activating the B_SEG_.

### RK neurons that induce wing expansion are environmentally sensitive

The ability of a stimulus to induce the full wing expansion program after a short delay increases with increasing stimulus duration (Fig. 2B, 1X). This would be expected if longer stimuli were more effective in countering the environmental inhibition posed by confinement. In this case, decreasing confinement—which has been previously shown to shorten the environmental search phase in a graded fashion (Peabody et al., 2009)—would be expected to increase the efficacy of shorter stimuli in inducing the wing expansion program. To test whether the efficacy of a stimulus of given duration is environmentally dependent, we placed flies in a cylindrical chamber identical to the minichamber, except five times longer. This environmental manipulation did not affect the immediate behavioral response to stimulation, but it did increase the effectiveness of shorter duration stimuli in inducing the wing expansion program. The half-maximum response to stimulation in the “5X” minichamber shifted to approximately 1.25 min of stimulation, compared with 2.5 min of stimulation in the minichamber (Fig. 2B, lower panel). The output pathway of the RK neurons that mediate the induction of the wing expansion program is thus sensitive to the magnitude of the environmental inhibition, indicating that these neurons are likely to play a role in the decision to expand rather than simply execute this decision based on the evaluation of environmental information by upstream circuitry. In contrast, the RK neurons responsible for the immediate behavioral effects of activation appear to have a strictly motor function and to act downstream of the wing expansion decision.

### Cholinergic RK neurons (RK^ChaT^) are required for inducing and sustaining wing expansion

The temporally separable responses to stimulation of the RK neurons indicates that they comprise at least two functionally distinct populations. To identify these populations, we used the Split Gal4 system (Luan et al., 2006b; Pfeiffer et al., 2010) to isolate subpopulations of RK neurons according to neurotransmitter phenotype. We drove UAS-dTrpA1 expression in cholinergic, glutamatergic, and GABAergic subsets of neurons and examined the behavioral responses to a five-minute activating stimulus (Fig. 3A). As expected, stimulation of positive control animals, which co-expressed rk-Gal4DBD and a ubiquitously expressed Tub-dVp16AD hemidriver, showed immediate induction of perching and abdominal contraction followed by activation of the full wing expansion program. Neither effect was observed in animals in which GABAergic RK neurons were activated, but activation of glutamatergic RK neurons elicited immediate abdominal contraction, albeit without perching or induction of the wing expansion program. In contrast, activation of cholinergic RK neurons lacked obvious immediate effects on abdominal contraction, but robustly induced delayed activation of the full wing expansion program (Video 3). In addition, selective stimulation of only non-cholinergic RK neurons (achieved by using rk^pan^-Gal4 to drive UAS-dTrpA1 in animals also carrying a ChaT-Gal80 transgene to block Gal4 activity in cholinergic neurons) resulted in the induction of only stimulus-delimited behavior without the induction of the full wing expansion program afterwards (Fig. 3B). These results indicate that glutamatergic and cholinergic subsets of RK neurons differentially regulate the motor versus the decision-making aspects of Bursicon action.

**Figure 3.**
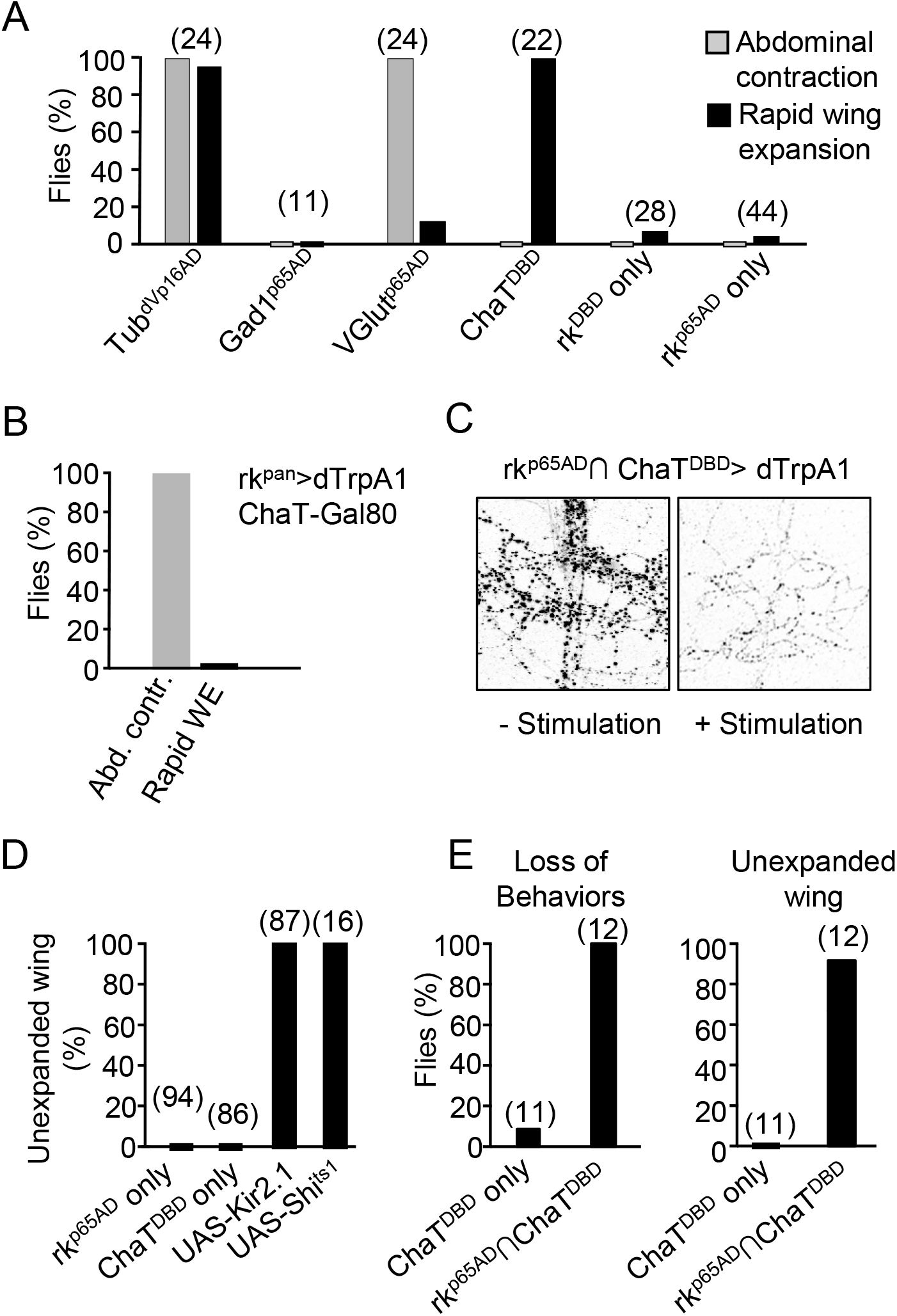
A cholinergic subset of RK neurons (RK^ChaT^) regulates Bursicon release and wing expansion via positive feedback. (A) Subsets of RK neurons were stimulated in newly eclosed, confined flies for 5 min and the effectiveness of the stimulus in inducing immediate abdominal contraction (gray) and rapid wing expansion (black) was measured in the indicated number of flies. GABAergic (Gad1^MI09277^-p65AD), glutamatergic (VGlut^MI04979^-p65AD), and cholinergic (ChaT^MI04508^-Gal4DBD) RK neuron subsets were targeted using the indicated Split Gal4 hemidrivers with corresponding rk^TGEM^-Gal4DBD or rk^TGEM^-p65AD hemidrivers. Negative control animals with only one RK hemidriver were also tested, in addition to positive control animals (rk^TGEM^-Gal4DBD∩Tub-dVp16AD) with expression throughout the RK expression pattern. n, in parentheses. (B) Selectively suppressing dTrpA1 expression in the cholinergic subset of RK neurons using the Gal4 inhibitor Gal80 (ChaT-Gal80) in combination with the rk^pan^-Gal4 driver blocks induction of rapid wing expansion (WE), while preserving the stimulus-delimited induction of abdominal contraction (Abd. Contr.). Bar graph shows the percentage of newly eclosed, confined flies that displayed each behavior in response to a 5 min stimulus (29°C). n=30. (C) Stimulation of RK^ChaT^ neurons is accompanied by Bursicon release from the B_SEG_. Images show anti-Bursα immunostaining of B_SEG_ projections in T2 of the VNC. No Stimulation: newly eclosed fly. With Stimulation: 90 min post-eclosion (fly confined and stimulated for 5 min immediately after eclosion). (D) Suppressing the RK^ChaT^ neurons by inhibiting either excitability (UAS-Kir2.1) or synaptic transmission (UAS-Shi^ts1^) completely blocks wing expansion. Flies eclosed without confinement into either a large food vial (UAS-Kir2.1) or a water-jacketed cuvette (UAS-Shi^ts1^) in which the temperature could be altered by changing the surrounding water temperature. In the latter case, synaptic block was accomplished by shifting the water temperature from 18°C to 31°C immediately after eclosion. Flies were observed for 4 hr after eclosion. (E) Acutely suppressing the RK^ChaT^ neurons after initiation of wing expansion terminates the execution of expansional behaviors (left panel) and blocks wing expansion (right panel). Control flies (ChaT^MI04508^-Gal4^DBD^>Shi^ts1^) or experimental flies (rk^TGEM^-p65AD∩ChaT^MI04508^-Gal4^DBD^>Shi^ts1^) were allowed to individually eclose into water-jacketed cuvettes, and after they perched and initiated abdominal contraction the temperature was shifted from 18°C to 31°C. All experimental flies ceased sustained abdominal contraction after the shift and most (11/12) had partially or completely unexpanded wings after 4 h at 31°C. In contrast, all control flies expanded their wings normally, although one such fly briefly ceased abdominal contraction.

Consistent with their role in biasing the wing expansion decision and inducing the wing expansion program, selective stimulation of the cholinergic subset of RK neurons (i.e. RK^ChaT^) also drives Bursicon release from B_SEG_ neuron fibers (Fig. 3C).

Complementing these results, we also found that constitutive suppression of RK^ChaT^ neurons using UAS-Kir2.1 inhibited wing expansion completely in flies that eclosed in the relatively unconfined environment of food vials (Fig. 3D). Similarly, acute suppression immediately after eclosion using UAS-Shi^ts1^ likewise inhibited expansion. These results are in contrast to those obtained when the B_SEG_ neurons are suppressed, in which case many unconfined flies succeed in expanding by grooming their wings open (Luan et al., 2012). The failure of flies in which the RK^ChaT^ neurons are suppressed to initiate expansion behaviors of any kind indicates that they are an essential component of the circuitry that drives wing expansion under all environmental conditions and are likely situated hierarchically above the B_SEG_.

To determine whether the RK^ChaT^ neurons are essential not only for initiating wing expansion behaviors, but also for sustaining them, we used UAS-Shi^ts1^ to acutely inhibit these neurons in flies that had already perched and initiated expansion, as indicated by fully flexed abdomens. All of the 12 experimental flies tested reverted to the unflexed state within several minutes and although most of them displayed several brief bouts of subsequent abdominal contraction (3.9 ± 4.0 bouts/30 min, stdv; n=12), it was never sustained and only one fly managed to fully expand its wings (Fig. 3E; Video 4). For almost all experimental flies (11/12), the wings remained unexpanded. In contrast, 100% of control animals (n=11) expanded their wings within the 30 min observation period and only one exhibited brief loss of abdominal flexion in response to the temperature shift.

### Signaling in the B_SEG_-RK^ChaT^ positive feedback loop

The above results are consistent with a model in which the B_SEG_ and RK^ChaT^ neurons participate in a positive feedback loop to drive wing expansion. In this model, Bursicon released from the B_SEG_ activates RK^ChaT^ neurons, which signal back to the B_SEG_ to promote further Bursicon release. To determine whether Bursicon signaling is required for RK^ChaT^ neuron induction of the wing expansion program, we selectively downregulated RK activity in RK^ChaT^ neurons. To do so, we used a membrane-tethered Bursicon construct, UAS-CFP-tBur-β-α (hereafter, UAS-tBur) known to desensitize RK *in vitro* and to disrupt Bursicon signaling in targeted cells *in vivo* (Harwood et al., 2014). When targeted to the RK^ChaT^ neurons, UAS-tBur causes wing expansion failure in 6% of flies allowed to expand in a large vial, a failure rate that increases approximately 10-fold when flies are confined in the minichamber (Fig. 4A). Bursicon signaling to the RK^ChaT^ neurons is thus particularly important for promoting the decision to expand under conditions of environmental inhibition.

**Figure 4.**
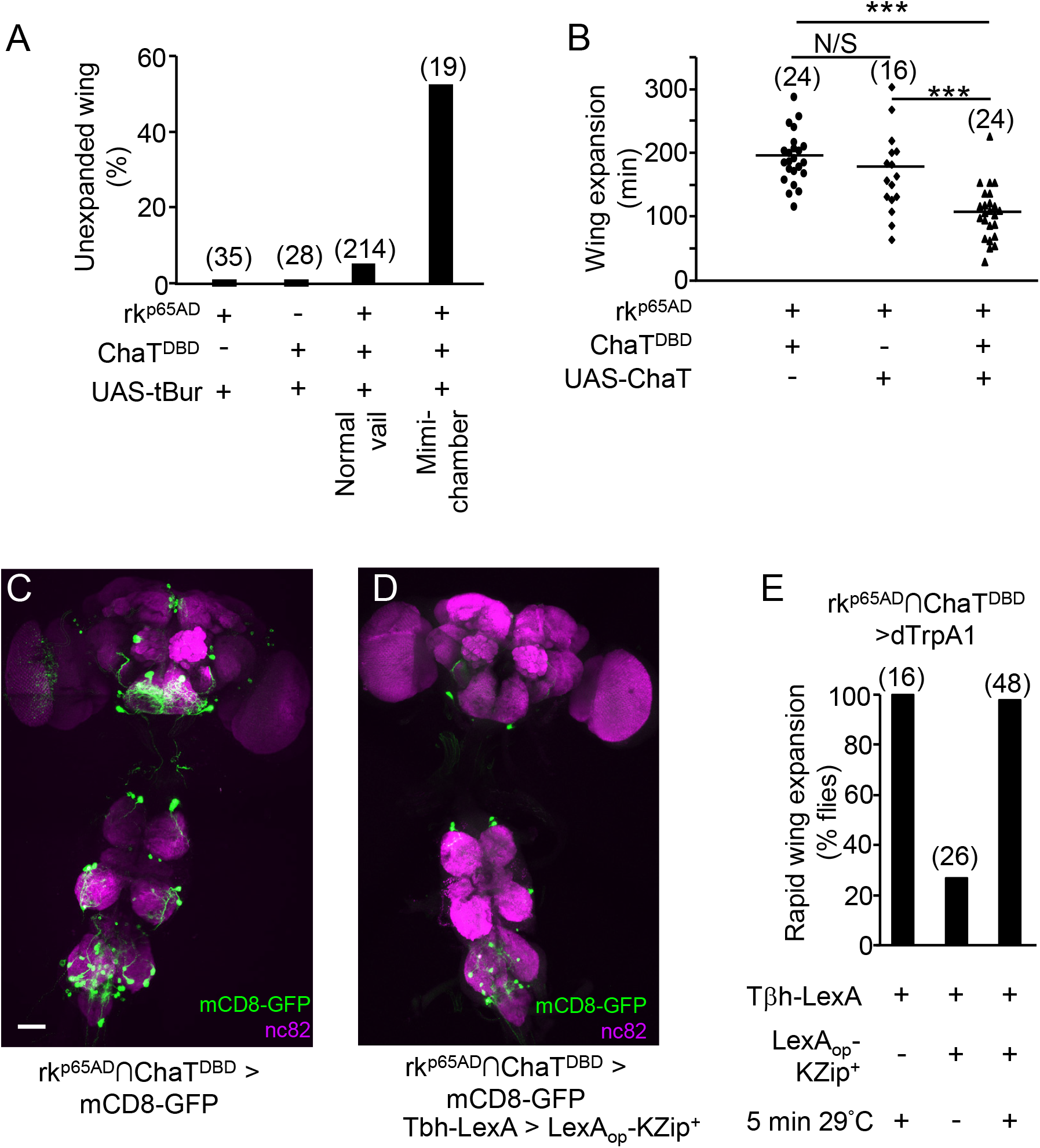
Characterization of RK^ChaT^ neurons: Bursicon and Ach signaling. (A) Constitutively expressing a membrane tethered Bursicon construct, UAS-tBur, in RK^ChaT^ neurons to desensitize RK receptors substantially blocks wing expansion in confined flies eclosing into a minichamber. Flies eclosing into a food vial are significantly less impaired, as are control flies lacking either of the rk^TGEM^-p65AD or ChaT^MI04508^-Gal4^DBD^ hemidrivers. n, in parentheses. (B) Flies over-expressing Choline Acetyltransferase (ChaT) in RK^ChaT^ neurons expand their wings significantly faster than control flies lacking either of UAS-ChaT or the ChaT^MI04508^-Gal4^DBD^ hemidriver. Scatter plots for each genotype show the expansion times of all newly eclosed flies in the minichamber at 25°C. Symbols as in Fig. 1A. n, in parentheses. (C) Confocal image showing RK^ChaT^ neurons visualized by UAS-mCD8-GFP reporter expression (green), with neuropil stained with nc82 antibody (magenta). (D) The complement of RK^ChaT^ neurons (UAS-mCD8-GFP; green) is substantially diminished when Split Gal4 activity is suppressed by LexA_op_-KZip^+^ expression in the subset of cholinergic neurons that lie within the pattern of a Tβh-LexA driver. nc82 antibody; magenta. (E) Activation of the RK^ChaT^ neurons induces rapid wing expansion even when the Split Gal4 inhibitor KZip^+^ is expressed in the Tβh-LexA pattern, as in (D).

Evidence that acetylcholine is likewise important in mediating communication between the RK^ChaT^ and B_SEG_ neurons comes from experiments in which a UAS-ChaT transgene was expressed in the RK^ChaT^ neurons to artificially enhance cholinergic neurotransmission. Flies expressing UAS-ChaT in the RK^ChaT^ neurons showed significantly advanced wing expansion in the minichamber when compared with controls, consistent with a facilitatory role for RK^ChaT^-derived Ach in wing expansion (Fig. 4B). However, efforts to assess synaptic connectivity between the RK^ChaT^ and B_SEG_ neurons using both anatomical and functional methods provided only negative evidence, suggesting that RK^ChaT^ neurons likely signal through one or more intermediaries in interacting with the B_SEG_.

Apart from feedback signals from the RK^ChaT^ neurons, the B_SEG_ are also expected to be interoceptive targets of ETH. While Bursicon-expressing neurons are known to express the A-isoform of the ETHR (ETHRA) at the pupal stage (Diao et al., 2016; Kim et al., 2006) this has not previously been confirmed for the adult stage. We therefore examined the overlap between ETHRA and Bursicon expression in pharate adults. As shown in Fig. 4—figure supplement 1, we find that adult Bursicon-expressing neurons also express ETHRA.

### Identification of RK^ChaT^ neurons that provide positive feedback to the B_SEG_

To characterize the population of RK^ChaT^ neurons, we examined the expression pattern of rk^p65AD^∩ChaT^DBD^ > mCD8-GFP flies (Fig. 4C). The expression pattern included approximately 30 neurons in the brain and over twice this number in the ventral nerve cord. We were able to further narrow this population by taking advantage of a fortuitous feature of Trojan exon insertions into the MiMIC^MI04010^ site in the Tyrosine β-Hydroxylase (Tβh) gene. Lines made with such insertions unexpectedly display extensive expression in cholinergic neurons (data not shown). A Tβh-LexA driver expressing the Split Gal4 repressor, LexA_op_-KZip^+^ (Dolan et al., ^2^01^7^) substantially diminished the number of neurons in the RK^ChaT^ pattern (Fig. 4D), but left induction of the wing expansion motor program completely intact when combined with rk^p65AD^∩ChaT^DBD^ > UAS-dTrpA1 (Fig. 4E). The reduced pattern of labeling consisted primarily of neurons in the ventral nerve cord and only two prominent neurons in the central brain. Interestingly, the latter neurons are in close proximity to the B_SEG_, but their identity remains to be determined.

### Glutamatergic RK neurons facilitate motor output

As described above, activation of the glutamatergic complement of RK neurons (RK^VGlut^) promptly induces abdominal contraction. The consequent elevation in internal pressure was evident from the fact that the wings of most flies (n=23/24) expanded partially during the period of activation, though in almost all cases they quickly refolded after activation. The induced abdominal contraction itself was somewhat unusual in that it only intermittently displayed the characteristic downward flexion seen normally during wing expansion, though the abdomens of all flies were clearly elongated. To determine whether this unusual phenotype was the result of the animals continuing to walk, we tethered flies while activating the RK^VGlut^ neurons. This allowed us to simultaneously assay for the induction of proboscis extension (PE) and cibarial pumping, two other expansional motor patterns which are difficult to observe in the minichamber (Fig. 5A). All tethered flies exhibited both abdominal elongation and flexion upon activation of RK^VGlut^ neurons, and all also extended their proboscises, though in no case was concomitant cibarial pumping observed (n=9). Activation of RK^VGlut^ neurons is thus sufficient to immediately evoke some, but not all of the motor patterns associated with wing expansion. Our preliminary evidence indicates that other populations of RK neurons, including a subpopulation of cholinergic neurons distinct from those that induce wing expansion after a short delay, may mediate other behavioral components of wing expansion, but these remain to be characterized.

**Figure 5.**
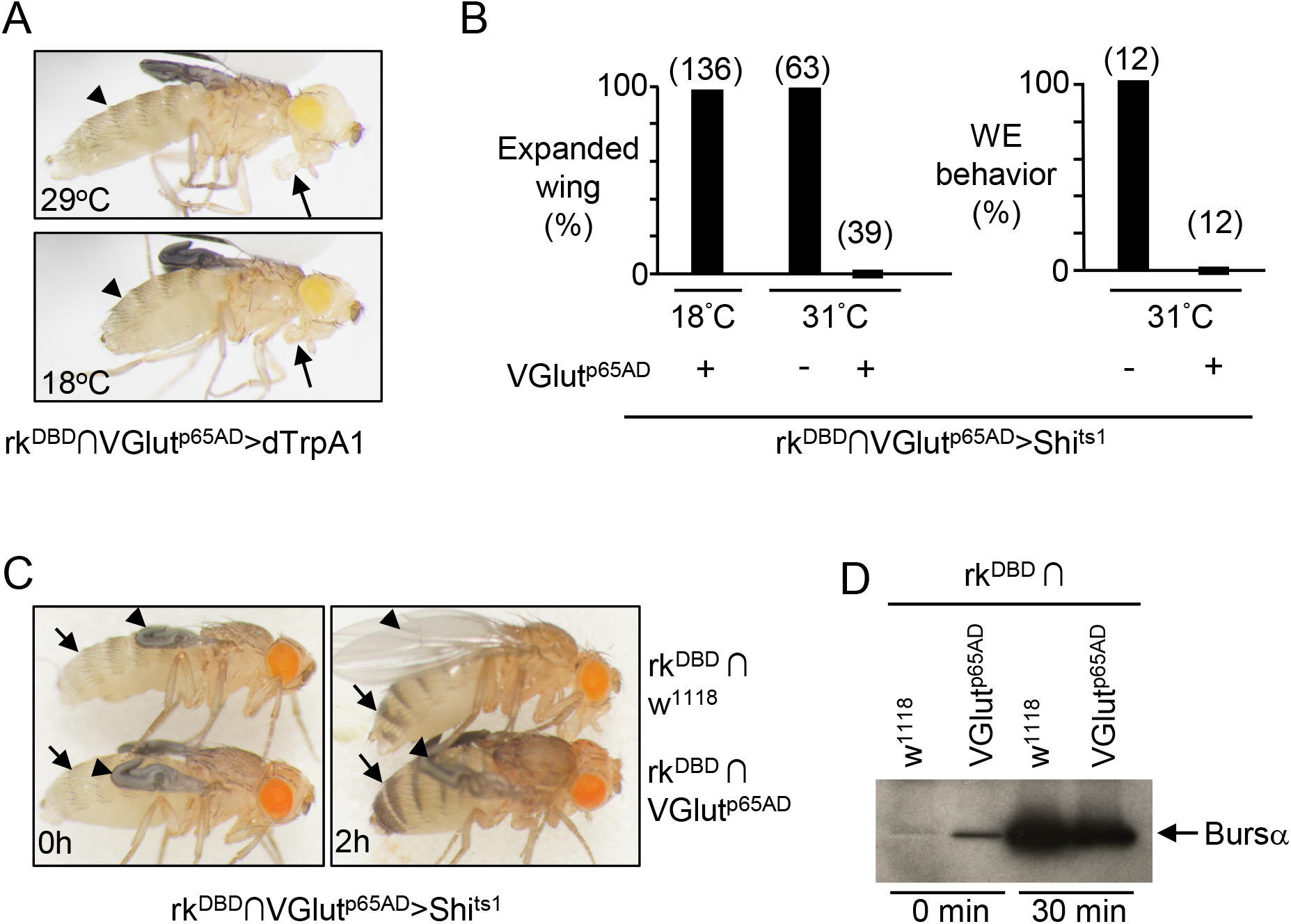
RK^VGlut^ neurons mediate wing expansion behaviors. (A) Stimulation of RK^VGlut^ neurons in a tethered fly using dTrpA1 immediately induces abdominal contraction (arrowheads) and PE (arrows). Fly is shown before (bottom) and during (top) stimulation. (B) Suppression of RK^VGlut^ neurons after eclosion using UAS-Shi^ts1^ blocks both wing expansion (left) and initiation of wing expansion behavior (right). Control flies in which RK^VGlut^ neurons are not suppressed because they remain at the restricted temperature or lack the VGlut^p65AD^ transgene expand normally. (**C-D**) Suppression of RK^VGlut^ neurons after eclosion does not block cuticle tanning or Bursicon release into the hemolymph from the B_AG_. (**C**) Normal tanning (or pigmenting) of the abdomen (arrows) is seen both in controls (top fly) and animals in which RK^VGlut^ neurons are suppressed (bottom fly) within 2 h after eclosion. Unlike control flies, which expand their wings (arrowheads) during this time, animals with suppressed RK^VGlut^ neurons fail to expand. (**D**) Bursicon titers in the hemolymph rise significantly within 30 min after eclosion (0 min) in both control animals (rk^DBD^∩w^1118^> Shi^ts1^) and animals in which the RK^VGlut^ neurons are suppressed (rk^DBD^∩VGlut^p65AD^>Shi^ts1^), as determined by Western blot using an anti-Bursα antibody.

To determine whether the RK^VGlut^ neurons are necessary for the execution of these motor patterns, we examined the effects of constitutively inhibiting their activity using UAS-Kir2.1. We find that this manipulation is lethal at adult ecdysis, with all animals failing to eclose despite showing preparatory motor activity, such as leg twitching. To obviate this lethality, we used UAS-Shi^ts1^. This manipulation completely prevents wing expansion in flies placed in cuvettes, an environment that permits normal expansion in control flies (Fig. 5B). Suppressed flies do not execute environmental search and often groom, but they show no evidence of abdominal contraction or PE. Despite the absence of wing expansion motor patterns, the flies do tan normally after 2 h (Fig. 5C) and examination of their hemolymph 30 min after eclosion indicates robust secretion of Bursicon from the B_AG_ (Fig. 5D). We conclude that RK^VGlut^ neurons are required for abdominal contraction and PE at the time of wing expansion, but not for perching, cibarial pumping, or the neuroendocrine release of Bursicon.

### Identification of behaviorally relevant subsets of RK^VGlut^ neurons

To identify the RK^VGlut^ neurons, we used a UAS-mCD8-GFP reporter driven by the rk-Gal4DBD and VGlut-p65AD hemidrivers and examined CNS expression by confocal microscopy (Fig. 6A). The number of labeled neurons observed was substantially reduced relative to the total complement of RK neurons, with most cell bodies located in the subesophageal zone (SEZ) and the ventral nerve cord, particularly in the abdominal ganglia. Some somata in the SEZ were located in areas known to contain cell bodies of glutamatergic motor neurons innervating the proboscis, namely the dorsomedial anterior (DMA) and ventrolateral posterior (VLP) clusters. To determine whether the RK^VGlut^ neurons included these motor neurons, we backfilled axons innervating the proboscis using rhodamine dextran and examined the clusters of motor neuron somata in the SEZ. To facilitate backfilling, we used 2-3d old flies after verifying that PE was also induced in such flies upon activation of RK^VGlut^ neurons (data not shown). We found four motor neurons in each hemisphere double-labeled with UAS-mCD8-GFP, one in the DMA cluster (Fig. 6B), and three in the VLP cluster (Fig. 6C). To determine the identity of these RK^VGlut^ motor neurons, we examined UAS-mCD8-GFP expression in head wholemounts after counterstaining the proboscis muscles with phalloidin (Fig. 6D). Consistent with the number of identified proboscis motor neurons, we identified four proboscis muscles innervated by labeled axons: M4, M8, M9, and M12-1. Interestingly, we also observed axons that extended medially down the proboscis, but which turned near muscle 9 then traveled caudally along the esophageal musculature, where they ramified broadly. Further work will be required to determine the nature of the interaction between the latter axons and the esophageal muscles, but we tentatively conclude that the RK^VGlut^ neurons include not only motor neurons that regulate proboscis extension, but also neurons that regulate visceral muscles of the alimentary tract, and which may play a role in swallowing air.

**Figure 6.**
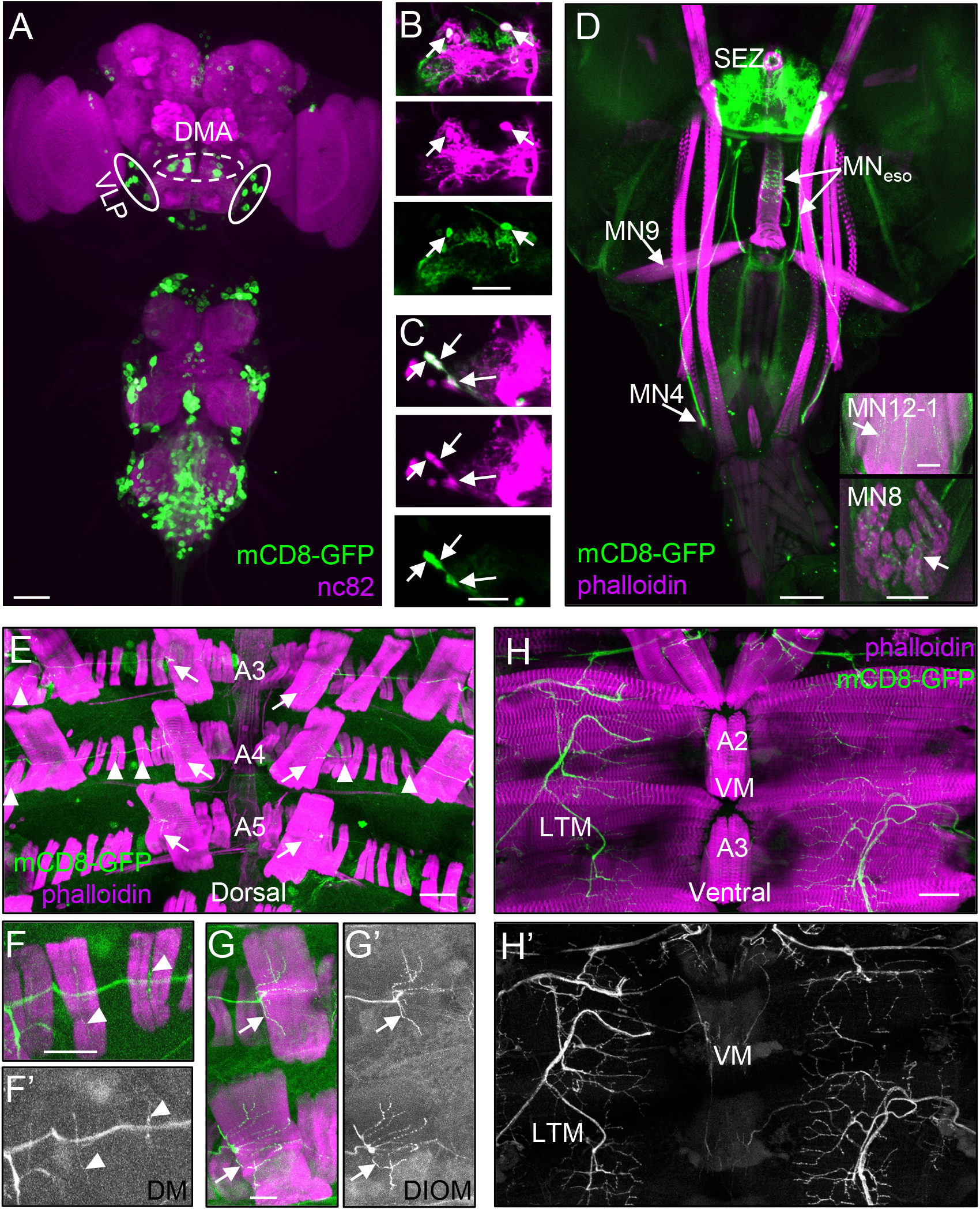
RK^VGlut^ motor neurons innervate proboscis and abdominal muscles. (**A**) RK^VGlut^ neurons in the pharate adult CNS visualized using UAS-mCD8-GFP (green). Regions that contain two clusters of proboscis motor neurons are indicated: the Dorsomedial Anterior (DMA, dotted oval) and the Ventrolateral Anterior (VLA, solid oval) clusters. Neuropil is counterstained with nc82 antibody (magenta). Scale bars: 50 μm and 25 μm in large and small images, respectively. (**B-C**) Four RK^VGlut^ neurons (arrows) are motor neurons within the (**B**) DMA and (**C**) VLP clusters of the SEZ. Proboscis motor neurons within these clusters were backfilled with rhodamine dextran (magenta, top and middle panels) and RK^VGlut^ neurons were visualized using mCD8-GFP (green, top and bottom panels). The DMA includes one, and the VLP three, bilateral pairs of RK^VGlut^ motor neurons. (**D**) Confocal micrograph of a head wholemount from a RK^VGlut^>mCD8-GFP fly showing muscles of the proboscis labeled with rhodamine-phalloidin (magenta) and motor axons labeled with mCD8-GFP (green). Terminals of motor axons (arrows) were identified on proboscis muscles 12 (MN12-1, top inset), 9 (MN9), 8 (MN8, bottom inset), 4 (MN4). Also projecting from the subesophageal zone (SEZ), were axons (MN_eso_) that ramified on the esophageal musculature. (Note, inset images are from a second head wholemount imaged from the opposite side.) (**E-H**) RK^VGlut^ motor neurons (mCD8-GFP, green or white) innervate dorsal and ventral muscles (rhodamine-phalloidin, magenta) of the adult abdomen. (**E**) Body wall fillet along the dorsal midline of segments A3-A5 shows motor axon terminals innervating numerous DMs (arrowheads) and the two DIOMs (arrows) flanking the midline. (**F**) Terminals on two DMs (arrowheads), showing labeled muscles and axons, or (**F’**) axons alone. (**G**) Terminals on two medial DIOMs (arrows) showing both muscles and axons, or terminals alone (**G**’). (**H**) Ventral midline body wall fillet of segments A2-A3 showing innervation of LTMs and VMs by RK^VGlut^ motor axons. (**H’**) Same as (H), showing axon terminal arbors only.

To determine whether the RK^VGlut^ neurons of the abdominal ganglia also include motor neurons that might participate in abdominal contraction, we examined fillet preparations of the abdominal musculature visualized with phalloidin from RK^VGlut^>UAS-mCD8-GFP animals. We identified muscles on both the dorsal and ventral body wall with GFP-positive terminals. On the dorsal side, these included numerous dorsal muscles (DM) in most segments examined (Fig. 6E, F, F’; arrowheads) as well as the two dorsal internal oblique (DIOM) muscles flanking the midline (Fig. 6G, G’; arrows). On the ventral side, the lateral tergoternal (LTM) and ventral (VM) muscles had axon terminals (Fig. 6H, H’). The DM, LTM, and VM muscles all persist after wing expansion, but the DIOM muscles have been previously shown to apoptose within 24 h of wing expansion and are thought to function specifically in adult ecdysis (Kimura and Truman, 1990).

## Discussion

Here we elucidate how a neuromodulator organizes a behavioral state by identifying and characterizing the targets of the B_SEG_, a pair of Bursicon-expressing neurons previously shown to positively regulate motor patterns used in wing expansion and to negatively regulate environmental inhibition of that process (Luan et al., 2012; Peabody et al., 2008). Our results confirm and clarify these dual roles and support a model in which cholinergic targets (RK^ChaT^ neurons), together with the B_SEG_, form a positive feedback loop that balances environmental contingencies and internal needs to promote the decision to expand, while glutamatergic RK neurons promote execution of the expansion decision (Figure 6—figure supplement 1). The circuit thus contains a positive feedback loop that promotes the transition from appetitive search to consummatory wing expansion and a feedforward pathway that targets the motor components of the consummatory action. Bursicon couples these two pathways, matching the motor requirements of a behavior to the motivation to execute it.

### The B_SEG_ act on diverse targets to command wing expansion

The circuit that regulates insect ecdysis is an example of what have been called “survival circuits,” which act at the interface between an animal’s essential needs and its environment (LeDoux, 2012; Sternson, 2013). A salient feature of many survival circuits—first noted by Hess in his seminal studies of the mammalian hypothalamus (Hess, 1949)–is that their electrical stimulation evokes robust and rapid expression of consummatory behaviors. For homeostatic circuits, the sites stimulated to evoke such behavior are often interoceptive neurons that signal using neuromodulators (for review see Augustine et al., 2020). In the case of the B_SEG_, neuromodulatory signaling by Bursicon is required for behavioral induction (Luan et al., 2012), which, as we show here, is not time-locked to B_SEG_ stimulation—typically starting only after a five minute stimulus—and is mimicked by acute elevation of the RK second messenger, cAMP, in RK neurons.

Our results reveal that Bursicon’s command of wing expansion is distributed to distinct types of downstream RK neurons. Short-latency RK^VGlut^ motor neurons encode separable motor components of wing expansion, while long-latency RK^ChaT^ neurons feed back to the B_SEG_. The induction of wing expansion following stimulation of RK^ChaT^ neurons likely reflects both to the time required for the RK^ChaT^ neurons to activate the B_SEG_ and the time required for the B_SEG_ to activate motor output. In addition to commanding activity in motor and feedback pathways, the B_SEG_ facilitate the neuroendocrine release of Bursicon from the B_AG_. This effect appears to be mediated by RK^ChaT^ neurons in that blocking these neurons blocks cuticle tanning, but further work will be required to determine the functionally relevant subgroup. Overall, the architecture of command described here—with short-latency components that mediate motor output and longer latency components that generate persistent internal states of activation—is similar to that described for several other survival circuits, including the circuits mediating rodent conspecific aggression (Falkner et al., 2020) and feeding (Chen et al., 2016), and fly grooming (Hampel et al., 2015) and courtship (Kennedy et al., 2014). Our data illustrate how the same mechanism that persistently maintains the internal state can be used to ensure sufficient drive to motor components to sustain behavior to completion.

### Positive feedback drives the transition from appetitive search to consummatory expansion

Neuromodulators contextualize behavior by biasing the probability of sensorimotor processes in response to environmental or physiological changes (Bargmann, 2012). Bursicon’s “command” of the wing expansion motor program exhibits particularly strong bias, derived from amplification of its own release via positive feedback. This use of positive feedback to potentiate hormone release has been observed in other systems (Ewer et al., 1997; Kingan et al., 1997; Moos et al., 1984). In the wing expansion circuit, positive feedback is used to ensure robust switching from search to expansion behavior and to sustain expansion behavior under normal conditions. Consistent with these observations, incremental suppression of all Bursicon-expressing neurons impairs the switch from search to expansion (Peabody et al., 2009) and full suppression of B_SEG_ function does not eliminate search behavior during the abnormal expansion observed in unconfined flies (Luan et al., 2012).

Once initiated, activity in the positive feedback loop between the B_SEG_ and RK^ChaT^ neurons is likely irreversible under natural conditions and represents the irrevocable commitment to expansion commonly observed. As first emphasized by early ethologists, the execution of many consummatory behaviors exhibits a similar all-or-nothing character (Craig, 1996; Lorenz, 1981), and positive feedback may be a common feature of survival circuits. It is noteworthy that the coupled feedback and feedforward pathways governing Bursicon action mirror those found elsewhere in the circuit architecture of ecdysis. In canonical models, ballistic ETH release is generated by a positive feedback loop with neurons that release EH (Ewer et al., 1997; Kingan et al., 1997), and ETH and EH exercise feedforward control of the motor program governing cuticle shedding (Ewer, 2007; Truman, 2005; Zitnan and Adams, 2012). In adult flies this motor program drives eclosion and subsequent environmental search and the factors that promote it (i.e. ETH and EH) both appear to signal forward to the B_SEG_ to promote expansion. The mechanisms by which they do so are poorly understood, but Bursicon-expressing neurons have been shown to respond to ETH exposure with delayed-onset Ca^++^ activity (Diao et al., 2017; Kim et al., 2015; Kim et al., 2006) and they are also potential targets of EH since ablation of EH-expressing neurons substantially blocks wing expansion (McNabb et al., 1997). The motor programs for appetitive search and consummatory expansion are thus regulated by sequential cycles of positive feedback coupled by feedforward signaling (Figure 6—figure supplement 1).

### Environmental regulation of positive feedback via the RK^ChaT^ neurons

The irreversible transitions promoted by positive feedback are both a benefit and a risk, since premature transitions are deleterious. Our evidence indicates that the RK^ChaT^ neurons play a critical role in preventing premature activity in the feedback loop and in matching behavioral output to environmental conditions. As indicated (Figure 6—figure supplement 1), these neurons are likely sensitive to conditions of confinement and require Bursicon signaling to overcome its effects. How Bursicon acts on the RK^ChaT^ neurons to relieve their sensitivity to environmental inhibition remains to be determined, but its action appears to be both cumulative and long-lasting. Partially suppressing Bursicon-expressing neurons slows the initiation of wing expansion (Peabody et al., 2009) and activating the B_SEG_ abrogates inhibition, but only after exceeding a threshold. Both of these observations are expected if Bursicon action is cumulative. Furthermore, if the B_SEG_ are briefly activated prior to eclosion, environmental inhibition of wing expansion is relieved for up to 45 min (Peabody and White, 2013) indicating the persistent effects of activation. This latter effect is reminiscent of the persistent effects of NPY on rodent feeding following stimulation of the hypothalamic AGRP neurons (Chen et al., 2019) and exemplifies what may prove to be a common role of neuromodulators in establishing the persistent internal states observed in numerous survival circuits (Anderson, 2016; Kennedy et al., 2014).

How are the B_SEG_ normally activated? We hypothesize that ETH initiates low-level Bursicon release from the B_SEG_ that does not activate downstream motor targets, but which incrementally affects—and eventually supersedes—environmental inhibition of the RK^ChaT^ neurons. This would explain why flies under conditions of continual confinement—such as in the minichamber—eventually decide to expand. In general, this model provides a heuristic framework for further characterizing the wing expansion network. The absence of identifiable anatomical or functional connections between the RK^ChaT^ and B_SEG_ neurons strongly suggests that other neurons are likely to participate in the positive feedback loop, and these may provide additional mechanisms for regulating its function. The circuitry that governs environmental search also remains to be identified, though, as indicated above, it is likely to be modulated by ETH and EH and thus be part of the broader ecdysis survival circuit. Understanding the interactions of the search circuit with components of the feedback loop will be critical for a complete understanding of the transition from appetitive to consummatory behavior in the search-or-expand decision.

### Neuroeconomics of the wing expansion decision

The search-or-expand decision of *Drosophila* lacks the complexity of decisions mediated by many survival circuits, such as those in mammals that regulate feeding, drinking, and salt appetite. These circuits are additionally regulated by learned cues, negative-feedback, and anticipatory-feedforward termination signals (Augustine et al., 2020; Sternson and Eiselt, 2017). They may also recruit the generalized mechanisms of decision-making that are used for performing learned tasks, such as those studied in non-human primates (Calhoun and Hayden, 2015; Pearson et al., 2014). It is interesting to consider the decision-making mechanisms described here in this broader context.

A critical adaptive feature of the wing expansion decision circuit is the positive feedback loop, which supplies the motivational drive to expand and encodes (via Bursicon) the motor consequences of the expansion decision. This tight coupling of drive to action ensures that the motor patterns governing wing expansion are immediately available for execution, should the decision to expand be made, but it also implies that the representation of the action—in its most reduced form, i.e. Bursicon—petitions for its own selection. Such coupling of decision and execution are a natural feature of reflexive behaviors, such as escape responses, in which dedicated neural determinants—sometimes single neurons—make and execute a behavioral decision (Edwards et al., 1999; Korn and Faber, 2005; von Reyn et al., 2014). However, it is also likely to operate in the higher-order, distributed decisions of primates where cognitive representations of possible actions are thought to compete for execution (Cisek, 2007; Pezzulo and Cisek, 2016). In the latter case, competition is biased by the performance costs of the competing actions, but little is known about the mechanisms that might couple such costs to the action. This problem is solved naturally in the example of the wing expansion circuit, which bundles action costs together with a description of the action and the motivation for executing it into B_SEG_ activity.

A critical difference between cognitive decisions and the type of decision made by the wing expansion circuit is that the former involve learned or improvised actions, the assembly and costs of which must be calculated using information derived by experience. In contrast, the behaviors underlying wing expansion, and the consummatory actions of survival circuits in general, are hard-wired. This means that their assembly and the costs of executing—or failing to execute— them under given environmental circumstances must come pre-calculated as genetically encoded biases in the circuitry. Numerous studies of basic behavioral decisions—for example, continued foraging under risk of predation—indicate that natural selection encodes valuations into the decision circuitry of animal brains to optimize survival or reproduction (Houston and McFarland, 1980; Houston and McNamara, 2014; Perry and Pianka, 1997). In the case of wing expansion, the valuations that guide the choice are presumably reflected in the weights of parameters that affect B_SEG_ activity, including the strength of inputs from hormones such as ETH and recurrent signals from the RK^ChaT^ neurons. This hypothesis remains to be tested, but the example of wing expansion suggests that the bundling of action costs and composition into one representation that then competes for execution might also be a strategy used by animal brains for making cognitive decisions in the service of survival.

## Author Contributions

Conceptualization, B.H.W., and F. D.; Methodology, F. D. and B.H.W.; Resources, N.C.P.; Investigation, F.D.; Writing, Review & Editing, B.H.W. and F.D.; Visualization, F.D. and B.H.W.; Supervision, B.H.W.; Funding Acquisition, B.H.W.

## Acknowledgements

This work was supported by the Intramural Research Program of the National Institute of Mental Health (ZIAMH002800, BHW). We further thank the Bloomington Drosophila Stock Center (NIH P40OD018537) for many of the fly stocks used in this study. Our very sincere thanks go to Tom Talbot and George Dold of the NIMH Section on Instrumentation Core Facility for custom design and construction of the manifold used to rapidly switch the temperatures of flies in up to five, individually housed, water-jacketed cuvettes, and the LED chambers used to optogenetically activate flies expressing UAS-euPACα. UAS-dTrpA1, UAS-CFP-tBurβ-α; and UAS-euPACα flies, were the kind gifts of P. Garity, Alan Kopin, and Martin Schwärzel.

The authors declare no conflicts of interest.

## Videos

**Video 1. Stimulation of RK neurons induces wing expansion.** Activating RK-expressing neurons using UAS-dTrpA1 at 29°C induces wing expansion. Negative control (top): w^1118^>dTrpA1; Experimental fly (bottom): rk^pan^>dTrpA1. Video speed: 40X.

**Video 2. Stimulating RK neurons for 2.5 min induces time-locked wing expansion behavior followed (after a delay) by the entire wing expansion program in almost half of flies.** A rk^pan^> dTrpA1 fly (top) responds to a 2.5 min temperature shift to 29°C with time-locked wing expansion behavior followed by stimulus-decoupled wing expansion after return to 18°C. Neither effect is observed in a w^1118^>dTrpA1 control fly (bottom). Video speed: 45X.

**Video 3**. **Briefly stimulating RK^ChaT^ neurons has no immediate behavioral effect, but elicits wing expansion after a short delay.** A 5 min temperature shift to 29°C has no immediate behavioral effect on experimental rk^p65AD^∩ChaT^DBD^ >dTrpA1 flies (+) in the minichamber, but several minutes after return to 18°C these flies initiate normal wing expansion. In contrast, rk^p65AD^ >dTrpA1 control flies (-) search throughout. Video speed: 40X.

**Video 4. RK^ChaT^ neurons are essential for maintaining wing expansion behaviors.** A temperature shift from 18°C to 31°C blocks synaptic activity of RK^ChaT^ neurons in rk^p65AD^∩ChaT^DBD^ > Shi^ts1^ flies. This manipulation terminates the wing expansion motor program in flies that have initiated it (note flexed abdomen and perching at video onset) and causes resumption of search behavior. Video speed: 40X.

## Supplemental Information

**Figure 1—figure supplement 1:**
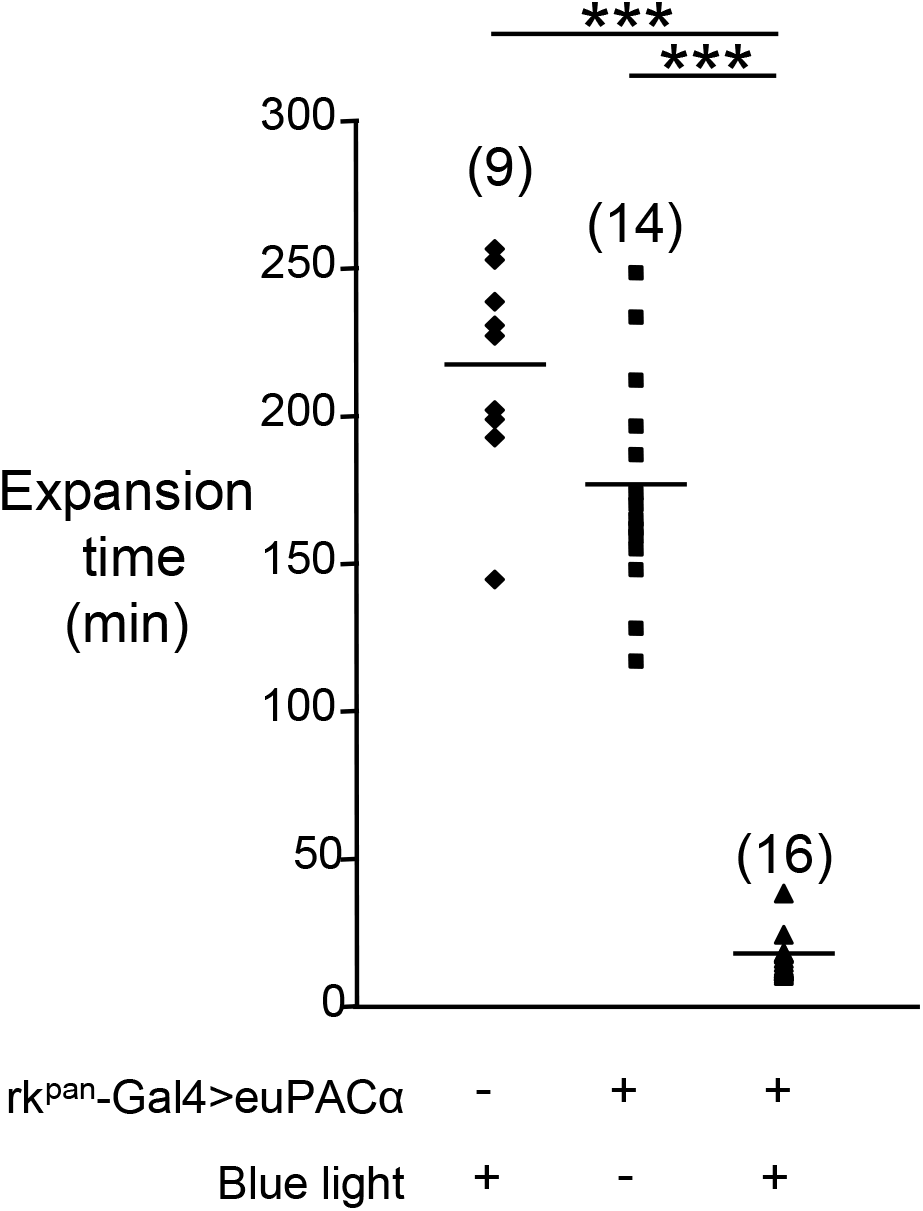
Photoactivation of adenyl cyclase activity in RK neurons induces rapid wing expansion. Photoactivation of the light-activated adenyl cyclase, UAS-euPACα, induces rapid wing expansion in newly eclosed, confined flies when expressed in RK neurons under the control of the rk^pan^-Gal4. Wing expansion times were measured in the minichamber after exposure to blue (450 nm) light for 2.5 min. Control flies lacked either the rk^pan^-Gal4 or UAS-euPACα transgenes and exhibited wing expansion times that were significantly delayed relative to experimental flies. Wing expansion times of individual flies are indicated, bars indicate the mean. n, in parentheses. ***, p<0.005 by Students’ t-test.

**Figure 1—figure supplement 2:**
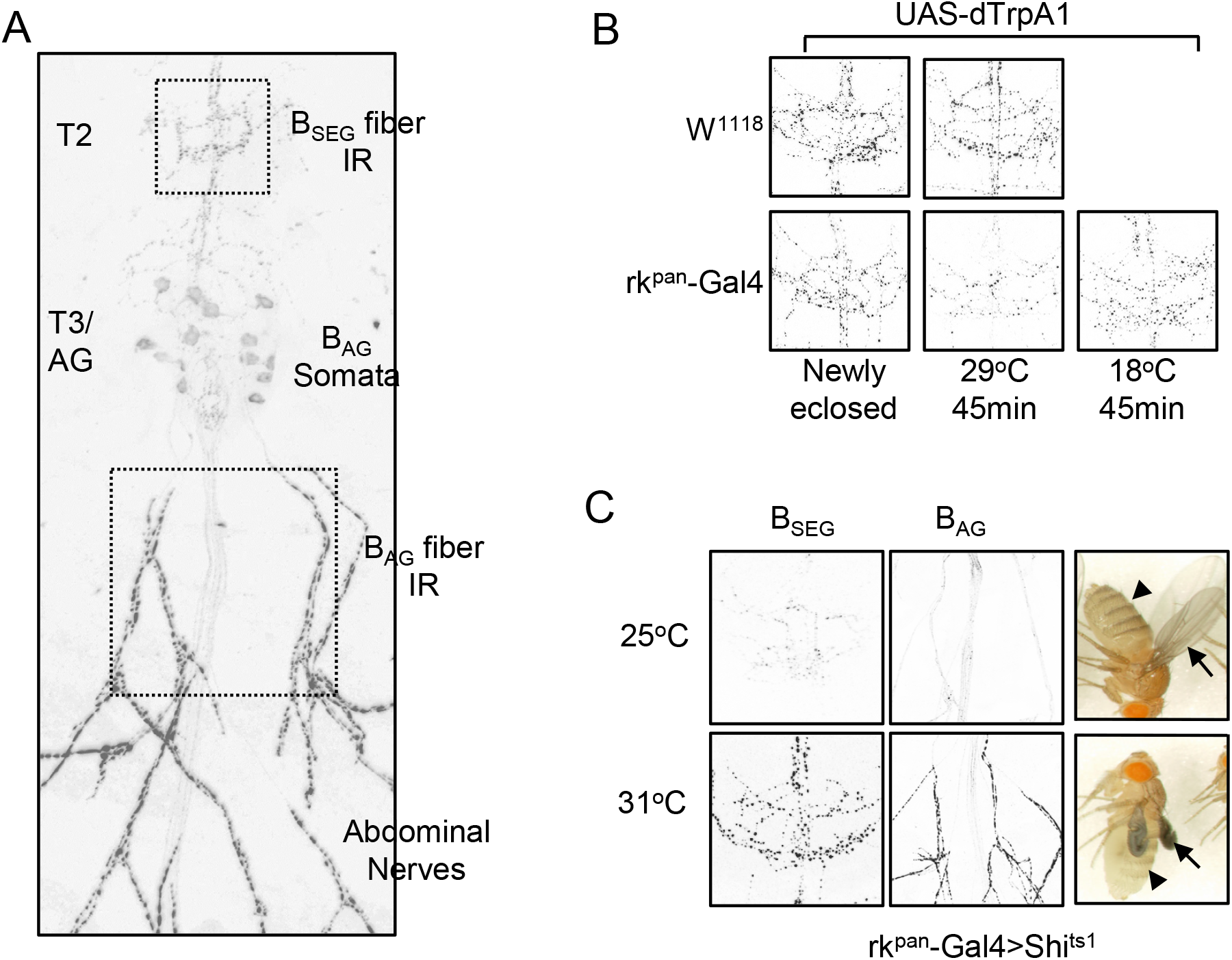
RK neuron stimulation causes Bursicon secretion. (A) Fillet preparation from a pharate adult showing anti-Bursicon immunoreactivity (IR) in the thoracic and abdominal ganglia (T2, T3, and AG) and the abdominal nerves exiting the ventral nerve cord (VNC). The Burs-IR in the abdominal nerves is due to the presence of B_AG_ fibers, while that in T2 reflects staining of B_SEG_ fibers. B_SEG_ Bursicon content is measured by quantifying the Burs-IR in a 200 x 200 pixel box framing T2 (upper box). B_AG_ Bursicon content is similarly measured from a 400 x 400 pixel box framing the exit point of the terminal abdominal nerves of the VNC (lower box). (B) RK neuron stimulation (45 min) causes Bursicon secretion from the B_SEG_. Representative images of Burs-IR in B_SEG_ fibers of the VNC in control (w^1118^>dTrpA1, upper panels) and experimental (rk^pan^-Gal4>dTrpA1, lower panels) animals. (C) Suppression of synaptic transmission in RK neurons of unconfined flies using UAS-Shi^ts1^ blocks wing expansion and Bursicon release from both the B_SEG_ and B_AG_. Neither B_SEG_ (left panels) nor B_AG_ (middle panels) Burs-IR is reduced in animals subjected to 1 hr RK neuron suppression (31°C), but both are reduced in animals spared from suppression (25°C), demonstrating that synaptic block of RK neurons inhibits Bursicon secretion. All animals in which Bursicon secretion is inhibited also fail to expand their wings (right panels, arrows) and to tan their cuticles (arrowheads).

**Figure 1—figure supplement 3:**
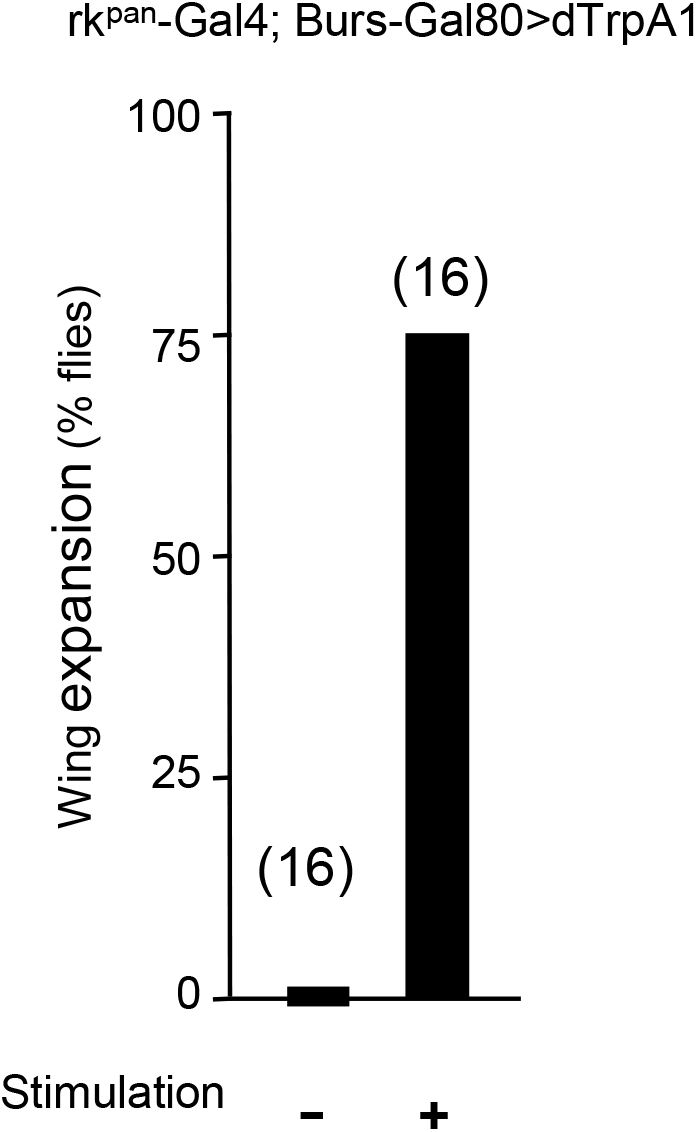
RK is not expressed in Bursicon-expressing neurons. Bar graph shows the percentage of newly eclosed, confined flies that respond to 5 min stimulation of the RK neurons by expanding their wings when UAS-dTrpA1 expression is selectively blocked in Bursicon-expressing neurons using a Gal80 transgene (Burs-Gal80). The stimulus induced wing expansion in 75% of flies (12/16), similar to the result of stimulating RK neurons in the absence of Burs-Gal80, as in Fig. 2B. Unstimulated control flies of the same genotype failed to expand rapidly.

**Figure 4—figure supplement 1:**
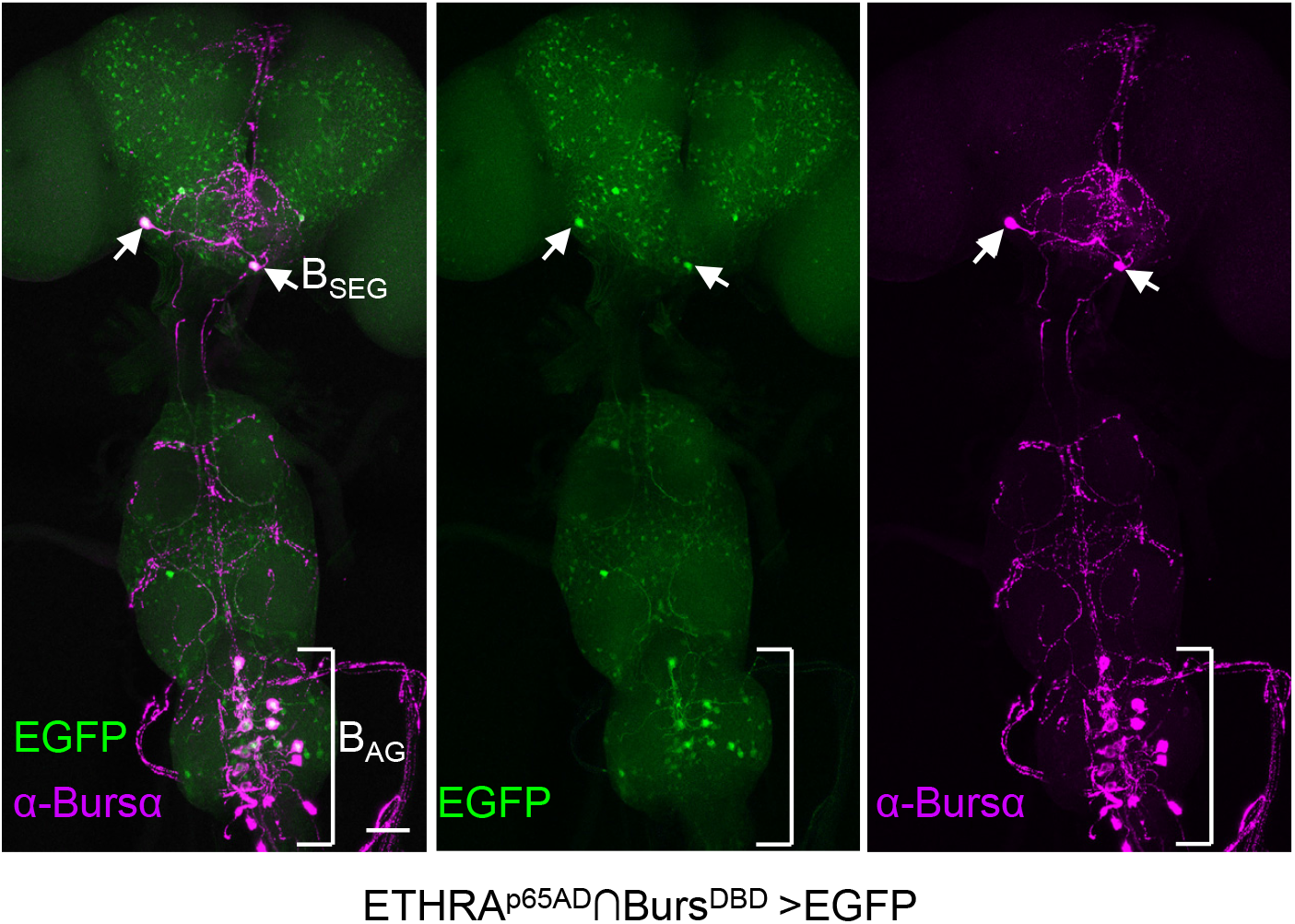
**ETH targets the B_SEG_** CNS wholemount prepared from a newly-eclosed animal expressing UAS-EGFP under the control of ETHRA^p65AD^∩Burs^Gal4DBD^ to target neurons that express both Bursicon and the A-isoform of the ETH receptor. The preparation was labeled with an anti-GFP antibody (green; left, middle) and immunostained with anti-Bursα antibody (magenta; left, right) to visualize the two populations of Bursicon-expressing neurons, the B_SEG_ (arrows) and B_AG_ (bracket). The merged images (left) reveal co-labeling of the B_SEG_, as well as a subset of the B_AG_ in anterior segments of the VNC. The image is a maximum projection of a confocal Z-stack.

**Figure 6—figure supplement 1:**
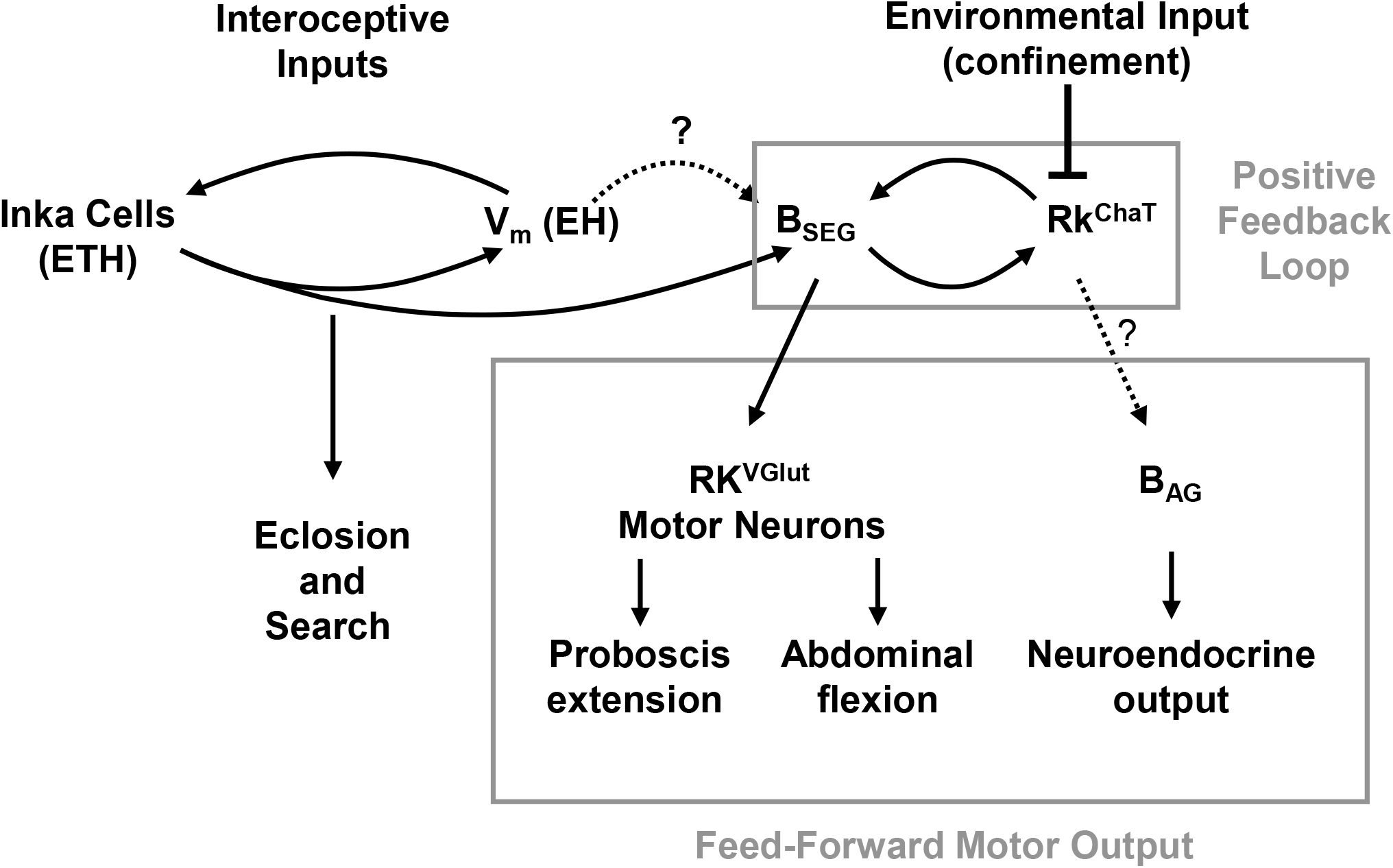
Organization of the decision circuit for wing expansion. Wing expansion under conditions of confinement is critically dependent on activity within a positive feedback loop formed between the B_SEG_ and RK^ChaT^ neurons. The previously observed sensitivity of wing expansion to environmental perturbations when the B_SEG_ are silenced, and the findings described here that the RK^ChaT^ neurons are required for wing expansion under all environmental circumstances and that their ability to drive wing expansion scales with the degree of confinement suggests that the RK^ChaT^ neurons are targets of environmentally-dependent inhibition. Bursicon released from the B_SEG_ is proposed to mitigate this inhibition onto the RK^ChaT^ neurons, which, once activated, drive activity of the B_SEG_ to promote ballistic release of Bursicon. Bursicon feeds forward to modulate RK^VGlut^ neurons that govern distinct wing expansion motor patterns. Bursicon also stimulates release of Bursicon from the B_AG_ via RK^ChaT^ neurons, though not necessarily the same population that participates in the feedback loop. B_SEG_ activity is thought to be positively regulated by the hormones EH and ETH, the release of which is similarly governed by a positive feedback loop and which mediate eclosion and search behaviors. In this model, the architecture of the ecdysis circuit thus generates sequential behavioral phases by coupling positive feedback loops that feed forward to promote distinct components of the sequence.

## Materials and Methods

### Key Resources Table

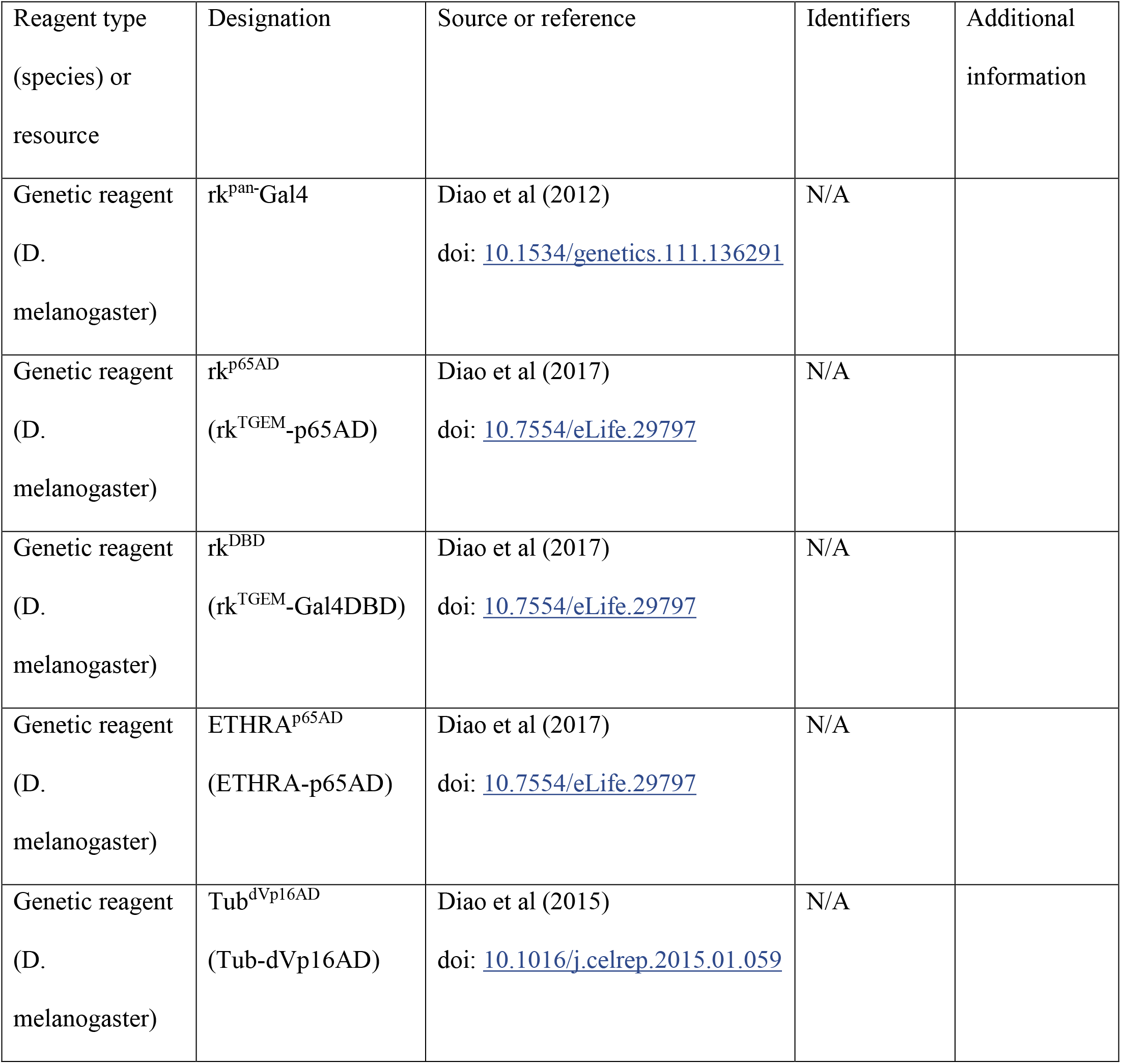
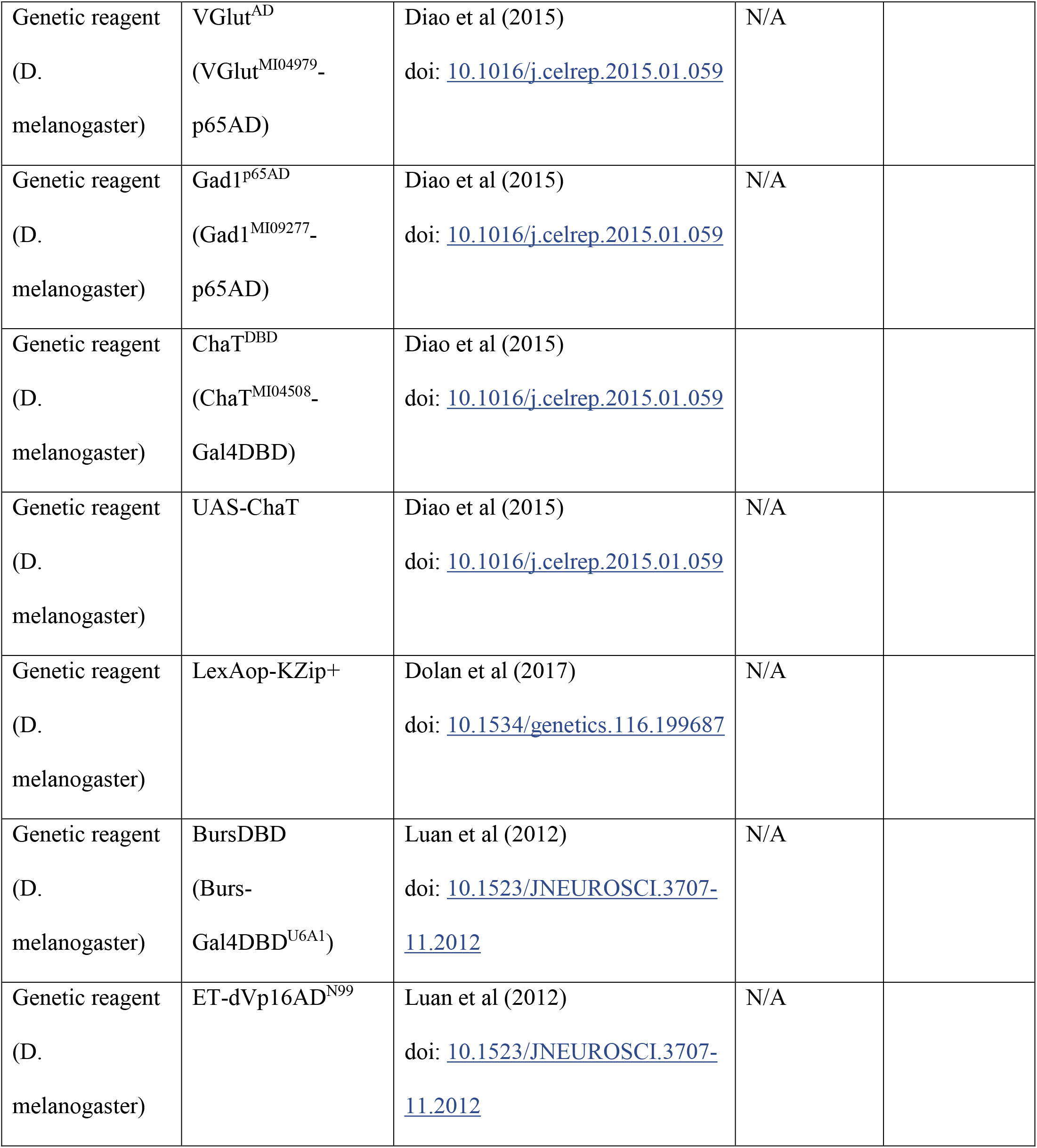
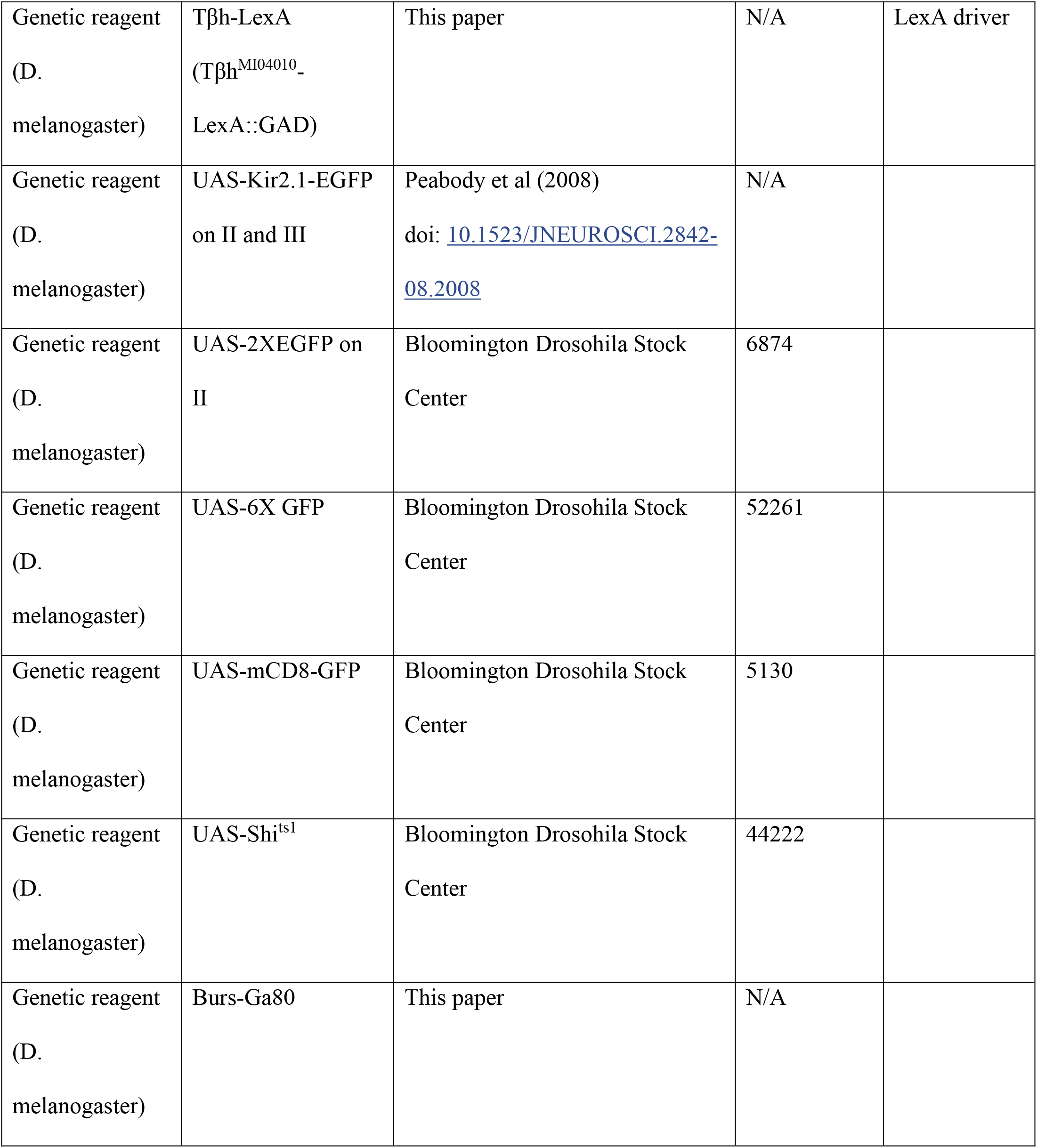
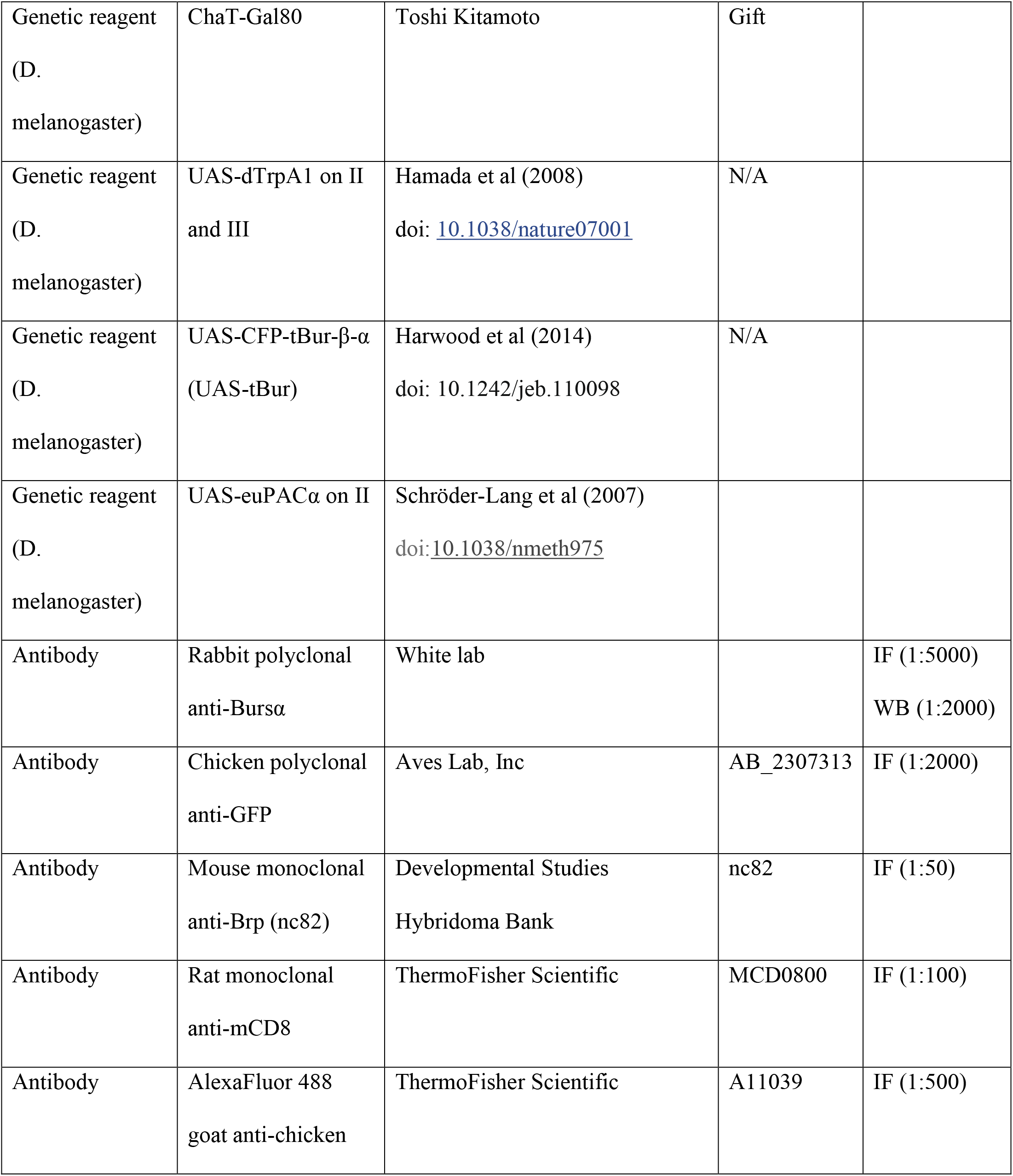
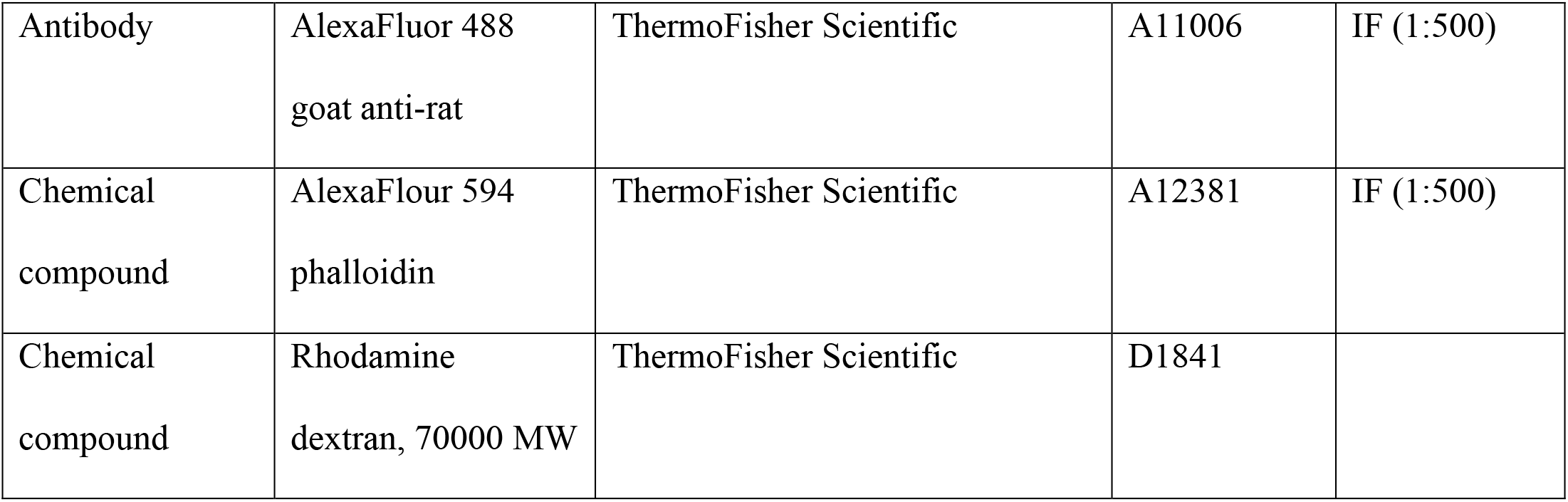

### Reagents

All PCR primers were synthesized by Integrated DNA Technologies, Inc (Coralville, IA). All other reagents are as listed in the Key Resources Table.

### Fly Stocks and Husbandry

Vinegar flies of the species *Drosophila* melanogaster were used in this study. Flies were raised on cornmeal-molasses-yeast medium and housed at 25°C and 65% humidity. Fly stocks described in previous publications (with alternative designations in parentheses) include: rk^pan^-Gal4 (Diao and White, 2012); rk^p65AD^ (rk^TGEM^-p65AD) and rk^DBD^ (rk^TGEM^-Gal4DBD), ETHRA^p65AD^ (ETHRA-p65AD) (Diao et al., 2017); Tub^dVp16AD^ (Tub-dVp16AD), VGlut^AD^ (VGlut^MI04979^– p65AD), Gad1^p65AD^ (Gad1^MI09277^–p65AD), ChaT^DBD^ (ChaT^MI04508^–Gal4^DBD^), UAS-ChaT (Diao et al., 2015); and LexA_op_-KZip^+^ (Dolan et al., 2017). The Burs^DBD^ (Burs-Gal4DBD^U6A1^) hemidriver and Split Gal4 B_SEG_ driver (i.e. ET-dVp16AD^N99^ ∩ Burs-Gal4DBD^U6A1^) are described in (Luan et al., 2012). Tβh-LexA (Tβh^MI04010^-LexA::GAD) line was generated by converting the indicated MiMIC line using the Trojan exon method as described in (Diao et al., 2015). All injections were made by Rainbow Transgenic Flies, Inc (Camarillo, CA). To make the Burs-Gal80 fly lines, the 252 bp Bursα promotor was amplified from the Burs-Gal4 construct (Peabody et al., 2008) and cloned into pCR-Blunt-II-TOPO (ThermoFisher Scientific, Inc) to make a “TOPO-Burs” intermediate. The Gal80 gene was then amplified from CCAP-Gal80 (Luan et al., 2006a) and cloned into the Not I/Xba I sites of TOPO-Burs, to make “TOPO-Burs-Gal80.” Spe I/Xba I double digests were then used to subclone the Burs-Gal80 DNA fragment into pCaSpR4 to make pCaSpR4-Burs-Gal80. Plasmid DNA from this construct was injected into w^1118^ embryos by Rainbow Transgenic Flies, Inc (CA) to make transgenic flies by p-element-mediated transformation.

The ChaT-Gal80, UAS-dTrpA1, UAS-CFP-tBur-β-α (UAS-tBur), and UAS-euPACα fly lines were the kind gifts of Drs. Toshi Kitamoto, Paul Garrity, Alan Kopin, and Martin Schwärzel, respectively. All other fly lines were obtained from the Bloomington Drosophila Stock Center at Indiana University.

### Videorecording and Behavioral Analysis

Flies were assayed for wing expansion and/or motor program execution, as previously described (Luan et al., 2012; Peabody et al., 2009). “Unconfined” environments were simulated using food vials or spectrophotometer cuvettes, which do not inhibit wing expansion. For environments with confined flies, glass “minichambers” (0.3 cm diameter x 0.7 cm length) or “5X” minchambers (0.3 cm diameter x 3.5 cm length) were used. For high resolution behavioral analysis of PE and cibarial pumping, flies were tethered to fine needles waxed to the thorax, given a cotton ball to hold with their legs and imaged directly under a microscope. Videorecording was accomplished with a Sony HDR-FX7 digital video camera. For flies in the minichamber or cuvette, we typically scored animals for success in wing expansion and/or initiation and completion of “abdominal contraction” (i.e. the persistent and readily observable extension and downward flexion of the abdomen). The time to wing expansion was defined as the time from eclosion until the expanded wings assumed their folded position over the abdomen. In cases where flies executed the wing expansion motor program, but failed to expand, expansion time was defined as the time from eclosion until the end of abdominal contraction (i.e. when the abdomen assumed its normal posture and was no longer extended and flexed). For experiments in which RK neurons (or subsets of RK neurons) were briefly activated, “rapid expansion” was defined as expansion in <120 min. (Note that expansion times are slow compared to those previously reported because, except for the time of stimulus delivery, the flies are at 18°C.) For experiments involving brief stimulation of the B_SEG_ neurons, “rapid expansion” was defined by the upper bound of expansion times for the 10 min stimulus. To assay for the induction of cibarial pumping and PE, newly-eclosed, tethered flies were videorecorded and observed. The time between heat application and cibarial pumping was scored and in some cases, air swallowing was confirmed by dissecting flies in glycerol and directly assaying for the presence or absence of air bubbles in the gut. For some genotypes, the induction of abdominal contraction and PE was also assayed in 2-3 d old tethered adults glued by the thorax to pipette tips.

### Activity Manipulations

#### Thermogenetic manipulations using UAS-dTrpA1 or UAS-Shi^ts1^

For experiments using confined flies, dTrpA1-mediated activation of neuronal activity was carried out in minichambers on a Peltier plate. Plate temperatures were adjusted to yield final chamber temperatures of 18°C (inactive) or 29°C. For activity suppression experiments using UAS-Shi^ts1^, animals were allowed to eclose in water-jacketed cuvettes placed on a custom manifold. The manifold has a plexiglass plate on one side that allows visual (i.e. video) access to the cuvettes. Water flow from one of two circulating water baths held at the target temperatures (18°C and 31°C) can be individually directed to the jacket of up to five cuvettes by solenoid switches. Typically, the cuvette temperature was shifted immediately after a fly was observed to eclose. Tethered flies were heated by adjusting their proximity to a Peltier plate held at the target temperature.

#### Optogenetic activation of RK-expressing neurons using euPACα

Most experiments using the light-activated adenylyl cyclase, UAS-euPACα, were carried out in a custom-designed aluminum chamber, painted black and mounted with 3 LUXEON Lumiled LEDs (Royal Blue = 450 ± 10 nm) on each of two sides. Newly eclosed dark-reared flies were collected into minichambers, which were put into a standard vial and inserted into the aluminum chamber. The flies in the minichambers were illuminated for 2.5 min and transferred onto a Peltier plate at 25°C for videorecording and counting wing expansion times.

#### Immunoblotting

Hemolymph collection and Western blotting were carried out as described in Peabody et al (2009). All animals were raised at 18°C and collected within 5 min of eclosion at room temperature. For stimulation experiments, minichambers containing experimental (rk^pan^-Gal4>UAS-dTrpA1) or control (w^1118^>UAS-dTrpA1 or rk^pan^-Gal4>W^1118^) flies were put on a Peltier plate for 10 min at 29°C, before the temperature was lowered to 18°C for 90 min. For the suppression experiment with RK^VGlut^, experimental (RK^VGlut^>Shi^ts1^) and control (rk^DBD^ only>Shi^ts1^) were collected into standard food vials and placed in a 31°C incubator for 30min. In all cases, hemolymph samples were pooled from ∼20 flies and 1 µL of hemolymph was loaded onto a 10% SDS-polyacrylamide gel for electrophoresis. Anti-Bursα antibody was used at 1:2000.

### Immunohistochemistry and Fluorescent labeling

#### Immunostaining and Image Acquisition

Excised nervous system whole mounts from pharate adults were dissected in Phosphate Buffered Saline (PBS), fixed in 4% paraformaldehyde/PBS for 20–30 min and then incubated in 4% paraformaldehyde/PBS plus 0.5% Triton X-100 for 15 min. Rabbit anti-Bursα (Luan et al., 2006b) was used at 1:5000 dilution and AlexaFluor 568 goat anti-rabbit secondary antibody (ThermoFisher Scientific.) was used at 1:500 dilution. Immunolabeled samples were mounted in Vectashield (Vector Laboratories, Burlingame, CA) prior to confocal imaging using a Nikon C-2 confocal microscope. *Z*-series were acquired in 1 μm increments using a 20× objective using 488 nm, and 543 nm laser emission lines for fluorophore excitation. Unless otherwise noted, the images shown are maximal projections of volume rendered *z*-stacks of confocal sections taken through the entire nervous system. For experiments in which proboscis motor neurons of RK^VGlut^>mCD8-GFP animals were backfilled, the tip of the proboscis of 2-3d old flies was removed near the maxillary palp with razor blade and a crystal of 70kD rhodamine dextran (ThermoFisher Scientific, Inc,) was placed on the cut. Flies were then incubated for 4h at 4°C prior to CNS dissection and immunostaining.

#### Immunolabeling fibers of the B_SEG_ and B_AG_

Flies were filleted and analyzed as described by Peabody et al. (2009). Briefly, animals were quickly anesthetized under CO_2_, immersed in 100% EtOH, then pinned out and filleted from the dorsal side in PBS. The head and internal organs were removed prior to fixation and staining. Z-series were acquired using a 20X objective, avoiding image planes containing the body wall. For quantitation of immunostaining, grayscale, volume-rendered images of the Z-stacks were inverted using Adobe Photoshop (Adobe Systems, Inc., San Jose, CA) and mean pixel values were calculated for a 400×400 pixel square centered over the abdominal nerves immediately after the exit point from the abdominal ganglion, or for a 50×50 pixel square centered over the rostral portion of the 2^nd^ thoracic segment, T2. Fibers in these regions were uniformly well-preserved in all preparations. Mean background pixel values (calculated for each image from a 200×200 pixel field outside of the imaged preparation) were subtracted to derive the “mean pixel intensities” used as a measure of bursicon immunoreactivity for each preparation. Statistical analysis of the means was performed by t-Test (two-sample assuming unequal variances).

#### Immunolabeling of axons innervating abdominal muscles

Pharate adults or newly eclosed flies were immersed in EtOH for 30 seconds and then transferred into PBS and pinned out by the head and the tip of the abdomen on either the dorsal or ventral side. The abdomen was then opened and pinned out, and the head, thorax and internal organs were removed. After fixation with 4% paraformaldehyde (PFA) for 30min and block for 2h at room temperature with 5% normal got serum in PBT (1X PBS with 0.1% Triton X-100), the filleted samples were incubated at 4°C for 2d with anti-mCD8 antibody (Thermo FisherScientific, Waltham, MA). Preparations were then washed in PBS 4X and incubated with secondary antibody (AlexaFluor 488 goat anti-rat) and AlexaFluor 594 phalloidin overnight at 4°C before washing and mounting.

#### Whole labeling of proboscis muscles and motor neurons

Whole heads were prepared as described by Schwarz et al (2017). Briefly, 2d-old RK^VGlut^>mCD8-GFP flies were fixed in 4% PFA with 0.1% Triton for 3h at 4°C with rotating. Heads were then severed and punctured with a minutien pin 10-12 times on the ventral side in PBS. Heads were washed with PBT 4 times and blocked with PBT/5% goat serum for 2 hr at room temperature. For immunostaining, heads were incubated at 4°C with rat anti-mCD8 antibody at 1:100 dilution for 7-8 days. After washing with PBT, the secondary antibody (AlexaFluor 488 goat anti-rat) and AlexaFluor 594 phalloidin were added for 4-5 days at 4°C with rotating. Both secondary antibody and phalloidin are diluted at 1:500. The confocal images were obtained with 20X objective.

#### Sample Sizes and Statistics

For all experiments, the sample sizes are indicated in the figure, figure legend or text. Significance testing was carried out using the Student t-test, unpaired 2-tail.

